# Differences in neurotoxic outcomes of organophosphorus pesticides revealed via multi-dimensional screening in adult and regenerating planarians

**DOI:** 10.1101/2022.06.20.496617

**Authors:** Danielle Ireland, Siqi Zhang, Veronica Bochenek, Jui-Hua Hsieh, Christina Rabeler, Zane Meyer, Eva-Maria S. Collins

**Author notes:** Correspondence: Eva-Maria S. Collins.

## Abstract

Organophosphorus pesticides (OPs) are a chemically diverse class of commonly used insecticides. Epidemiological studies suggest that low dose chronic prenatal and infant exposures can lead to life-long neurological damage and behavioral disorders. While inhibition of acetylcholinesterase (AChE) is the shared mechanism of acute OP neurotoxicity, OP-induced developmental neurotoxicity (DNT) can occur independently and/or in the absence of significant AChE inhibition, implying that OPs affect alternative targets. Moreover, different OPs can cause different adverse outcomes, suggesting that different OPs act through different mechanisms. These findings emphasize the importance of comparative studies of OP toxicity. Freshwater planarians are an invertebrate system that uniquely allows for automated, rapid and inexpensive testing of adult and developing organisms in parallel to differentiate neurotoxicity from DNT. Effects found only in regenerating planarians would be indicative of DNT, whereas shared effects may represent neurotoxicity. We leverage this unique feature of planarians to investigate potential differential effects of OPs on the adult and developing brain by performing a comparative screen to test 7 OPs (acephate, chlorpyrifos, dichlorvos, diazinon, malathion, parathion and profenofos) across 10 concentrations in quarter-log steps. Neurotoxicity was evaluated using a wide range of quantitative morphological and behavioral readouts. AChE activity was measured using an Ellman assay. The toxicological profiles of the 7 OPs differed across the OPs and between adult and regenerating planarians. Toxicological profiles were not correlated with levels of AChE inhibition. Twenty-two “mechanistic control compounds” known to target pathways suggested in the literature to be affected by OPs (cholinergic neurotransmission, serotonin neurotransmission, endocannabinoid system, cytoskeleton, adenyl cyclase and oxidative stress) and 2 negative controls were also screened. When compared with the mechanistic control compounds, the phenotypic profiles of the different OPs separated into distinct clusters. The phenotypic profiles of adult vs regenerating planarians exposed to the OPs clustered differently, suggesting some developmental-specific mechanisms. These results further support findings in other systems that OPs cause different adverse outcomes in the (developing) brain and build the foundation for future comparative studies focused on delineating the mechanisms of OP neurotoxicity in planarians.

## 1 Introduction

Organophosphorus pesticides (OPs) are among the most agriculturally important and common pesticides used today (EUROSTAT, 2016; Atwood and Paisley-Jones, 2017), especially in developing countries (Kaur and Singh, 2020). OPs kill pests by inhibiting the enzyme acetylcholinesterase (AChE) (Russom et al., 2014; Costa, 2018; Taylor, 2018), which is responsible for hydrolyzing the neurotransmitter acetylcholine (ACh). AChE inhibition causes cholinergic toxicity, ultimately manifesting as paralysis and death, in both insects and humans. This mechanism makes OPs effective pesticides, but because it acts the same way in humans as in insects, it is necessary to avoid human exposure to acutely toxic OP concentrations. This is of special concern in developing countries, such as India (Kaur and Singh, 2020) and South Africa (Razwiedani and Rautenbach, 2017), where accidental OP poisoning is prevalent. Moreover, growing evidence correlates chronic prenatal and infant exposure to subacute levels of OPs with life-long neurological damage and behavioral disorders (Rauh et al., 2011; Muñoz-Quezada et al., 2013; González-Alzaga et al., 2014; Shelton et al., 2014; Burke et al., 2017; Sagiv et al., 2021). These epidemiological studies are especially alarming given the environmental abundance of OPs.

Moreover, some OPs have been shown to affect secondary targets independent of and/or in the absence of significant AChE inhibition. OPs have been found to bind to human albumin (Peeples et al., 2005), and to rodent muscarinic ACh receptors (AChRs) (Howard and Pope, 2002; Lein and Fryer, 2005; Proskocil et al., 2010), demonstrating that OPs can directly interact with other proteins. OP-induced DNT in animals in the absence of significant AChE inhibition has been linked to a multitude of other secondary targets as well, depending on the system, OP, and exposure protocol (Dam et al., 2000; Slotkin, 2006; Slotkin et al., 2006b; Yang et al., 2008; Brown and Pearson, 2015; Mamczarz et al., 2016; Schmitt et al., 2019). Other proposed secondary targets include nicotinic AChRs, other esterases, and non-esterase, non-cholinergic targets such as serotonin receptors, cytoskeletal proteins, mitochondria, and glial cells (Pope, 1999; Guizzetti et al., 2005; Pope et al., 2005a; Slotkin et al., 2006b, 2017; Carr et al., 2014; Burke et al., 2017). The impact of all of these effects remains unclear, however, because it has been difficult to ascertain direct connections between molecular/cellular endpoints and brain function (behavioral) deficits.

Most mechanistic OP research has been focused on chlorpyrifos (CPF). Animal studies have shown that neurodevelopmental low-level exposure to CPF or its active metabolite, CPF-oxon (CPFO), can cause increased oxidative stress, cell death, and structural and functional neuronal deficits (Crumpton et al., 2000; Caughlan et al., 2004; Levin et al., 2004; Slotkin, 2004, 2006; Yang et al., 2011). CPF has been shown to bind to the microtubule associated motor protein kinesin (Gearhart et al., 2007) and CPFO to bind to tubulin and affect polymerization (Prendergast et al., 2007; Grigoryan et al., 2008; Jiang et al., 2010), implying that observed defects in axonal outgrowth and transport in animal studies may be a direct consequence of these interactions. The effects of developmental CPF exposure in animals have been found to be irreversible (Slotkin et al., 2001). These studies support epidemiological findings (e.g., (Rauh et al., 2011, 2012)) suggesting a causal link between developmental CPF exposure and long-term negative health effects. These data prompted the U.S. EPA to ban all food uses of CPF by 2022. Now other, less studied OPs are among the pesticides replacing CPF. Whether these OPs are indeed safer alternatives or also cause adverse effects on neurodevelopment is unclear. It is difficult to extrapolate the low dose toxicity profiles of other OPs from data on CPF, because OPs are structurally diverse with known pharmacokinetic (Pope, 1999; Jansen et al., 2009) and possible pharmacodynamic differences (Pope, 1999; Pope et al., 2005a; Terry, 2012). Comparative studies in rats have shown that different OPs damage the developing brain to varying extents, resulting in different adverse outcomes (Moser, 1995; Pope, 1999; Slotkin et al., 2006a; Richendrfer and Creton, 2015; Voorhees et al., 2016) and effects on developmental trajectories (Slotkin et al., 2006b), reinforcing the need to thoroughly evaluate individual OPs to better understand any potential compound-specific toxicity. Thus far, however, studies have largely been limited to either 1-4 compounds at a time (Slotkin et al., 2006a, 2006b, 2007; Yen et al., 2011; Richendrfer and Creton, 2015; Schmitt et al., 2019) or to studying acute and short term effects (Moser, 1995; Cole et al., 2004; Koenig et al., 2020).

To fill this data gap, we utilized high-throughput screening (HTS) in the asexual freshwater planarian *Dugesia japonica* to compare the toxicity profiles of 7 OPs (acephate, CPF, dichlorvos, diazinon, malathion, parathion, and profenofos). These OPs were chosen because of their environmental abundance, differences in chemical structures, and known potency in planarians from our previous work quantifying the *in vitro* inhibition rates of the respective oxons (Hagstrom et al., 2017a). Some of these OPs require metabolic activation by cytochromes P450 into their oxon form to inhibit AChE (CPF, parathion, diazinon, and malathion) and some can directly inhibit AChE (dichlorvos, profenofos, acephate). We have previously shown that exposure to diazinon, which does require bioactivation, results in significant AChE inhibition in both adult and regenerating planarians, suggesting that planarians can bioactivate OPs at all life stages (Hagstrom et al., 2018).

We have demonstrated that *D. japonica* is a unique and apt system for developmental neurotoxicology (Hagstrom et al., 2015, 2016, 2019; Zhang et al., 2019a, 2019b; Ireland et al., 2020). Neuro-regeneration is the sole form of neurodevelopment in this asexual species and shares fundamental processes with vertebrate neurodevelopment (Cebrià, 2007; Hagstrom et al., 2016; Ross et al., 2017). Thus, neurodevelopment can be induced by amputation, wherein the tail piece will regenerate a new brain within 12 days (Hagstrom et al., 2016). As intact and decapitated planarians are of similar size, adult and regenerating specimen can be tested in parallel with the same assays, providing the unique opportunity to directly identify effects specific to neurodevelopment. The planarian central nervous system, while morphologically simple, has considerable cellular and functional complexity (Cebrià, 2007; Ross et al., 2017). Planarians and mammals share key neurotransmitters (Ribeiro et al., 2005; Pagán, 2014), including ACh, which has been shown to regulate motor activity in *D. japonica* (Nishimura et al., 2010). Moreover, we identified 2 putative genes responsible for cholinesterase function in *D. japonica*, which were sensitive to OP inhibition and whose knockdown recapitulated some phenotypes of subacute OP exposure (Hagstrom et al., 2017a, 2018). Lastly, planarians have a variety of different quantifiable behaviors which can be assayed to assess neuronal functions. Importantly, several of these behaviors have been shown to be coordinated by distinct neuronal subpopulations (Nishimura et al., 2010; Inoue et al., 2014; Birkholz and Beane, 2017) allowing us to link functional adverse outcomes with cellular effects.

To begin to delineate the molecular mechanisms underlying OP toxicity, we compared the toxicological profiles of 7 OPs to chemicals with known modes of action (Table 1). These included cholinergic activators, such as carbamate AChE inhibitors (aldicarb and physostigmine) and nicotinic and muscarinic AChRs agonists (nicotine/anatoxin-a and muscarine/bethanechol, respectively). By testing both AChE inhibitors and the receptor agonists, we hoped to parse out effects specific to the nicotinic or muscarinic systems, especially since some OPs have been found to target the receptors directly (Howard and Pope, 2002; Smulders et al., 2004; Lein and Fryer, 2005; Proskocil et al., 2010). We also tested compounds known to affect alternative targets suggested in the literature to be affected by OPs. First, as cytoskeletal proteins such as actin and tubulin have been suggested to be direct targets of OPs (Jiang et al., 2010; Flaskos, 2012, 2014; Zarei et al., 2015), we tested actin polymerization inhibitors, cytochalasin D and latrunculin A, and anti-mitotic drugs (microtubules), nocodazole and colchicine. Second, fatty acid amide hydrolase (FAAH) has been shown to be inhibited by CPF leading to accumulation of the endocannabinoid anandamide and subsequent activation of the CB-1 receptor (Casida and Quistad, 2004; Liu et al., 2013; Carr et al., 2014). Thus, we characterized the toxicological effects of anandamide and the CB-1 receptor agonist WIN 55 212-2, which has previously been shown to affect planarian behavior (Buttarelli et al., 2002). We used mianserin, sertraline, buspirone, and fluoxetine to study potential disruption of the serotonergic system, which has been found to be sensitive to OP exposure during development (Aldridge et al., 2005; Slotkin et al., 2006b; Slotkin and Seidler, 2008), and LRE-1 and MDL-12,330A to study inhibition of adenyl cyclase, another proposed developmental OP target (Song et al., 1997; Slotkin and Seidler, 2008). Lastly, to test the effects of oxidative stress, which has been linked to OP DNT (Crumpton et al., 2000; Fortunato et al., 2006), we evaluated the effects of rotenone and L-buthionine sulfoxime.

**Table 1.** Chemical overview. . NA: not available.

| Chemical Name | CAS | DTXSID | Class/mode of action | Concentration range<sup>1</sup> | Supplier | Purity (%) |
| --- | --- | --- | --- | --- | --- | --- |
| Acephate | 30560-19-1 | DTXSID8023846 | OP | 1.78 – 316 | Sigma-Aldrich | 98 |
| Chlorpyrifos | 2921-88-2 | DTXSID4020458 | OP | 0.178 – 31.6 | Sigma-Aldrich | 100 |
| Diazinon | 333-41-5 | DTXSID9020407 | OP | 0.0316 – 31.6 | Sigma-Aldrich | 98 |
| Dichlorvos | 62-73-7 | DTXSID5020449 | OP | 0.00562 – 3.16 | Sigma-Aldrich | 98 |
| Malathion | 121-75-5 | DTXSID4020791 | OP | 0.316 – 56.2 | MP Biomedicals | 96 |
| Parathion | 56-38-2 | DTXSID7021100 | OP | 0.178 – 31.6 | Sigma-Aldrich | 100 |
| Profenofos | 41198-08-7 | DTXSID3032464 | OP | 0.0562 – 10 | Chem Service | 97 |
| Aldicarb | 116-06-3 | DTXSID0039223 | Carbamate AChE inhibitor | 3.16 – 316 | Sigma-Aldrich | 98 |
| Physostigmine | 57-47-6 | DTXSID3023471 | Carbamate AChE inhibitor | 0.1 -10 | Sigma-Aldrich | 99 |
| Anatoxin-A | 64285-06-9 | DTXSID50867064 | Nicotinic AChR agonist | 1 – 100 | Abcam | 98 |
| Nicotine | 54-11-5 | DTXSID1020930 | Nicotinic AChR agonist | 10 – 1000 | Sigma-Aldrich | 98 |
| Muscarine chloride | 2936-25-6 | DTXSID40861854 | Muscarinic AChR agonist | 1 – 100 | Sigma-Aldrich | 98 |
| Bethanechol chloride | 590-63-6 | DTXSID2022676 | Muscarinic AChR agonist | 3.16 – 316 | TCI America | 98 |
| Buspirone hydrochloride | 33386-08-2 | DTXSID1037193 | Serotonin (5-HT) 1 receptor agonist | 1 – 100 | Sigma-Aldrich | 99 |
| Mianserin hydrochloride | 21535-47-7 | DTXSID30944145 | Serotonin (5-HT) 1 receptor antagonist; Histamine H1-receptor agonist | 1 – 100 | Sigma-Aldrich | 98 |
| Fluoxetine hydrochloride | 56296-78-7 | DTXSID7020635 | Selective serotonin reuptake inhibitor | 1 – 100 | Sigma-Aldrich | 98 |
| Sertraline hydrochloride | 79559-97-0 | DTXSID1040243 | Selective serotonin reuptake inhibitor | 1 – 100 | Sigma-Aldrich | 98 |
| MDL-12,330A | 40297-09-4 | DTXSID10432999 | Adenyl cyclase inhibitor | 1 – 100 | Sigma-Aldrich | 98 |
| LRE-1 | 1252362-53-0 | NA | Adenyl cyclase inhibitor | 1 – 100 | Sigma-Aldrich | 98 |
| Colchicine | 64-86-8 | DTXSID5024845 | Disrupts microtubule polymerization | 3.16 – 316 | Acros Organics | 97 |
| Nocodazole | 31430-18-9 | DTXSID9031800 | Disrupts microtubule polymerization | 0.1 – 10 nM | Sigma-Aldrich | 98 |
| Cytochalasin D | 22144-77-0 | DTXSID8037099 | Disrupts actin polymerization | 0.316 – 31.6 | MP Biomedicals | 99 |
| Latrunculin A | 76343-93-6 | DTXSID90893488 | Disrupts actin polymerization | 0.316 – 31.6 nM | Sigma-Aldrich | 95 |
| Anandamide | 94421-68-8 | DTXSID301017453 | Endocannabinoid | 1 – 100 | Sigma-Aldrich | 97 |
| WIN 55 212-2 mesylate | 131543-23-2 | DTXSID50424974 | CB-1 receptor agonist | 0.1 – 10 | Sigma-Aldrich | 98 |
| L-buthionine sulfoxime | 83730-53-4 | DTXSID70894150 | Induces oxidative stress | 0.1 – 10 mM | Sigma-Aldrich | 97 |
| Rotenone | 83-79-4 | DTXSID6021248 | Induces oxidative stress | 3.16 – 316 nM | Sigma-Aldrich | 100 |
| L-ascorbic acid | 50-81-7 | DTXSID5020106 | Negative control | 1 – 100 | Alfa Aesar | 99 |
| D-glucitol | 50-70-4 | DTXSID5023588 | Negative control | 1 – 100 | Sigma-Aldrich | 99 |

<sup>1</sup>unless otherwise stated, concentrations are in $\mu\text{M}$

Using this comparative screening approach, we found differences in neurotoxicity and DNT for the 7 OPs in adult and regenerating planarians, respectively. Toxicological profiles were not correlated with levels of AChE inhibition. Using hierarchical clustering of the phenotypic profiles, we identified 6 clusters each in adult and regenerating planarians. The endpoints affected by the OPs and hierarchical clustering of OP phenotypes with those induced by the mechanistic control compounds differed between adult and regenerating planarians, suggesting that the planarian system can detect development-specific toxicity. For both worm types, we found a cluster that was indicative of cholinergic toxicity. However, certain OPs and concentrations were also found in the other clusters, implying effects on alternative targets. Together, our data show that planarian HTS can recapitulate the diverse toxicity profiles of OPs that have been observed in other systems and that these differences cannot be explained by levels of AChE inhibition. Cluster analysis suggests that OPs affect multiple targets in planarians and that the adverse outcomes differ depending on the developmental stage, emphasizing the need for more comparative OP studies to better understand the mechanisms by which OPs damage the nervous system.

## 2 Materials and Methods

### 2.1 Test animals

Freshwater planarians of the species *Dugesia japonica,* originally obtained from Shanghai University, China and cultivated in our lab for > 8 years, were used for all experiments. Planarians were stored in 1x Instant Ocean (IO, Blacksburg, VA) in Tupperware containers and kept at 20°C in a Panasonic refrigerated incubator in the dark. The animals were fed either organic freeze-dried chicken liver (either Mama Dog’s or Brave Beagle, both from Amazon, Seattle, WA) or fresh organic chicken or beef liver from a local butcher once a week. Their aquatic environment was cleaned twice a week following standard protocols (Dunkel et al., 2011). For all experiments, only fully regenerated worms which had not been fed within one week and which were found gliding normally in the container were used. Planarians were manually selected to fall within a certain range of sizes, with larger planarians used for amputation/regeneration experiments, such that the final sizes of adult and regenerating tails were similar. To induce development/regeneration, intact planarians were amputated on day 1 by cutting posterior to the auricles and anterior to the pharynx with an ethanol-sterilized razor blade. Exposure began within 3 hours of amputation.

### 2.2 Chemical preparation

Table 1 lists the chemicals used in this study. Two presumed negative control chemicals, D-glucitol and L-ascorbic acid, previously shown to not affect planarian behavior or morphology (Zhang et al., 2019a), were also screened. L-ascorbic acid was inactive in all endpoints as expected. D-glucitol was active in one locomotion-based outcome measure each in adult and regenerating planarians at the highest tested concentration (100 µM). Both of these outcomes were from new endpoints we had not previously evaluated. Thus, D-glucitol at 100 µM may not be an appropriate negative control for this system. Chemical stock solutions were prepared in 100% dimethyl sulfoxide (DMSO, Sigma-Aldrich, Saint Louis, MO) or milliQ water depending on solubility. All stock solutions were stored at −20°C. Each OP was tested at 10 concentrations over quarter-log steps. For all other chemicals, 5 concentrations were tested using semi-log steps. The highest concentrations were chosen, based on preliminary tests, to be at the threshold to cause lethality or overt systemic toxicity so that we could focus on sublethal behavioral effects. If lethality was not observed the highest soluble concentration was used. Diazinon and dichlorvos were found to have significant effects at the lowest tested concentrations in our preliminary analysis and thus were rescreened at 3 and 2 lower concentrations (quarter-log steps), respectively.

Chemical stock plates were prepared in 96-well plates (Genesee Scientific, San Diego, CA) by adding 200X stock solutions from the highest tested concentration to one well of the plate. Serial dilutions were then made in DMSO or IO water. The control well contained pure DMSO or IO water. In the final screening plates, either 0.5% DMSO, which has no effects on planarian morphology or behavior (Hagstrom et al., 2015), or IO water were used as solvent controls. Stock plates were sealed and stored at −20 °C for up to 3 months. For the OPs, the screening data comes from two separate screens due to a laboratory relocation. For all other chemicals, all concentrations were screened together in one of the two screens, with the chemicals split between the two locations. Colchicine was screened in duplicate to evaluate consistency of results between the two screens. The results between the two screens were consistent at low to medium concentrations, but we obtained a few hits (body shape at day 7 and day 12 and scrunching) at the highest concentration in one of the screens. These small differences in sensitivity may be due to changes in food source and availability, as previously discussed (Zhang et al., 2019a).

### 2.3 Screening plate setup

Each 48-well screening plate (Genesee Scientific) assayed 8 planarians in the solvent control (0.5% DMSO or IO water), and 8 planarians each per concentration of chemical (5 test concentrations per plate). Experiments were performed in at least triplicate (independent experiments performed on different days). Some chemical conditions and specific assays were repeated due to poor health in the control population or technical malfunction. The orientation of the concentrations in the plate was shifted down 2 rows in each replicate to control for edge effects (Zhang et al., 2019a). For each chemical and experiment, one plate containing adult (intact) planarians and one plate containing regenerating tails (2 plates total) were assayed. For the OPs, 2 plates of each worm type were screened to cover the 10 test concentrations.

Screening plates were prepared as described in (Zhang et al., 2019a) with one adult planarian or tail piece in each well of a 48-well plate containing 200 µl of the nominal concentration of test solution and sealed with ThermalSeal RTS seals (Excel Scientific, Victorville, CA). The plates were stored, without their lids, in stacks in the dark at room temperature when not being screened. Since we previously found that worms that underwent asexual reproduction (fission) produced challenges in our automated data analysis pipeline (Zhang et al., 2019b) and because planarian fission is suppressed when disturbed (Malinowski et al., 2017), the plates were gently agitated by hand once every 1-2 days when not being screened to discourage fission. Prepared plates were only moved to the screening platform when screened at day 7 and day 12.

### 2.4 Screening platform

Screening was performed on an expanded version of the planarian screening platform described in (Zhang et al., 2019a; Ireland et al., 2020). Briefly, this platform consists of a commercial robotic microplate handler (Hudson Robotics, Springfield Township, NJ) and multiple cameras and assay stations, which are computer controlled. Outcomes measures were obtained from studying planarian behavior on the assay stations (phototaxis/locomotion/morphology, stickiness, thermotaxis, and noxious heat sensing/scrunching). We modified phototaxis from the assay described in (Zhang et al., 2019a; Ireland et al., 2020) to measure the planarians’ response to different wavelength (red, green, blue) light using RGB lights (DAYBETTER, Shenzhen, China). Planarians are insensitive to red, detect green with their eyes, and blue with their skin pigment and eyes (Brown et al., 1968; Paskin et al., 2014; Birkholz and Beane, 2017; Shettigar et al., 2017). Therefore, using green and blue light stimuli allows us to discern between effects specific to the photoreceptors (green light) versus effects on extraocular perception through the skin. The phototaxis assay was performed as follows: First, to lower the variability of the animals’ background activity, planarians were allowed to acclimate for 3 min. After this acclimation period, the plate was imaged for 5 minutes: 1-min red light (1^st^ dark cycle), 1-min green light (light cycle), 2-min red light (2^nd^ dark cycle), 1-min blue light (light cycle). The temporal reaction of the planarians to the different light periods was quantified by calculating the average speed in every 30 second interval using center of mass tracking. Morphology data was collected by either imaging planarians using 4 cameras as described in (Zhang et al., 2019a) or by imaging with a single high resolution camera (Basler acA5472, Basler, Germany), with both imaging methods yielding the same resolution and number of frames per well. Identification of lethality and different abnormal body shape categories (Ireland et al., 2020) was performed manually by a reviewer who was blind to the chemical identities. Stickiness was conducted as described in (Ireland et al., 2020), and measures the number of worms that are stuck/unstuck when the plate is shaken at a fixed rotation per minute (RPM). Thermotaxis and scrunching were conducted as described in (Zhang et al., 2019a; Ireland et al., 2020), measuring the planarians’ response to low temperature gradients and noxious heat, respectively.

We have also added additional endpoints to quantify anxiety and locomotor bursts (Supplementary Figure 1). Anxiety was measured during the second dark phase of the phototaxis assay and calculated as time spent at well boundary / total time tracked. Higher anxiety scores represent less exploration as planarians prefer gliding along the container wall (Akiyama et al., 2015). We defined locomotor bursts as instances when a planarian accelerated from resting (speed <0.2 mm/s) to moving (speed >0.2 mm/s) and calculated the total cumulative number of locomotor bursts and the ratio of locomotor bursts in the blue period compared to the second dark period during phototaxis. Only blue light was used because planarians are most sensitive to short wavelengths (blue/ultraviolet) (Paskin et al., 2014; Shettigar et al., 2017) and thus their behavior changes are more robust in blue versus green light. These new endpoints were assayed on both day 7 and day 12. Data was primarily analyzed using MATLAB (MathWorks, Natick, MA). Data analysis was performed blinded with no chemical information provided. Chord diagrams were created using the circlize package (Gu et al., 2014) in R (R Core Team, 2016).

### 2.5 Benchmark concentrations (BMCs)

Benchmark concentrations (BMCs) were calculated for every outcome measure and chemical to quantify potency using the Rcurvep R package (Hsieh et al., 2019). Data for the different worm types (adult and regenerating) and for each day were treated independently. First, the outcome measures were transformed into an amenable format to allow for determination of directional, concentration-dependent responses. For all binary endpoints (lethality, body shape, stickiness, scrunching and eye regeneration), the incidence rates (number of planarians affected and total number of planarians) from the combined data from all replicates (n≥24) was used (Table 2). Normalizing by the vehicle control response was not necessary for lethality, body shape and eye regeneration as the control response rate was generally 0 (Supplementary Figure 2). For stickiness and scrunching, which had a more variable control response (Supplementary Figure 2), the experimental response was normalized by the incidence number of the respective in-plate vehicle controls. In some cases, this normalization led to negative incidence rates in the experimental responses. Because the Rcurvep package cannot handle negative incidence rates and these values are within the variability of the control populations, we only considered increases in abnormal activity. Thus, these negative incidence rates were set to 0. For continuous endpoints, the raw response of each individual planarian was normalized either by dividing by or subtracting by the median of vehicle control values for that plate (Table 3).

**Table 2.** Binary endpoints. The standard deviation (SD) of the vehicle controls and benchmark response (BMR) are compared for each endpoint on day 7(d7) and day 12 (d12), except for eye regeneration and scrunching which were only evaluated on d7 and d12, respectively.

| Endpoint | Description | Adult |  | Regenerating |  |
| --- | --- | --- | --- | --- | --- |
|  |  | SD (%) | BMR (%) | SD (%) | BMR (%) |
| Lethality | % dead | d7: 1<br>d12: 3 | d7: 10<br>d12: 20 | d7: 0<br>d12: 2 | d7: 10<br>d12: 15 |
| Body shape | % individuals with any abnormal body shape | d7: 2<br>d12: 6 | d7: 20<br>d12: 25 | d7: 6<br>d12: 6 | d7: 30<br>d12: 25 |
| Stickiness | % stuck individuals | d7: 26<br>d12: 22 | d7: 50<br>d12: 50 | d7: 18<br>d12: 20 | d7: 50<br>d12: 50 |
| Eye regeneration | % individuals with abnormally regenerated eyes | ---- |  | d7:12 | d7:55 |
| Scrunching | % non-scrunching planarians | d12: 11 | d12: 25 | d12: 13 | d12: 50 |

**Table 3.**
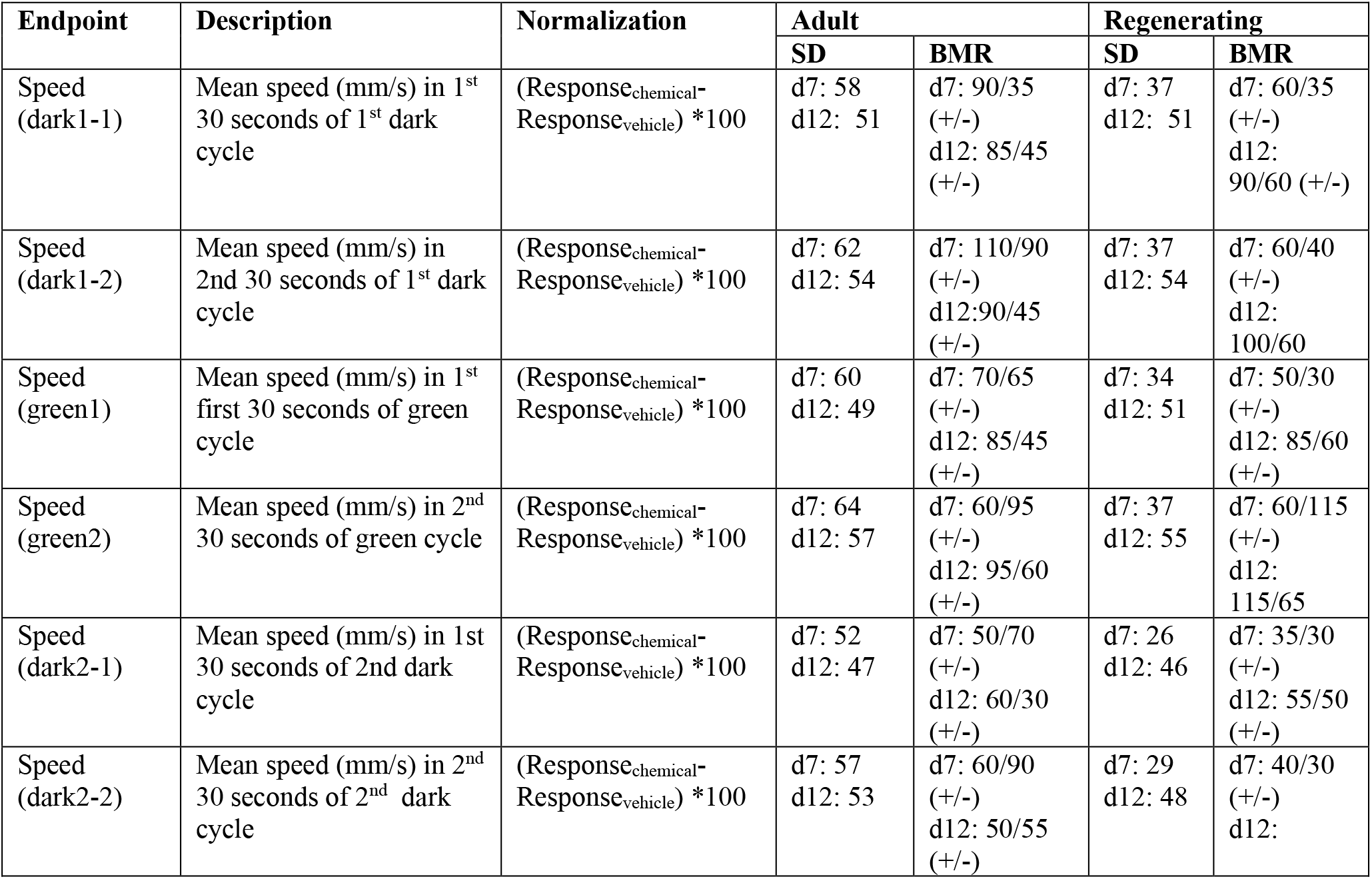
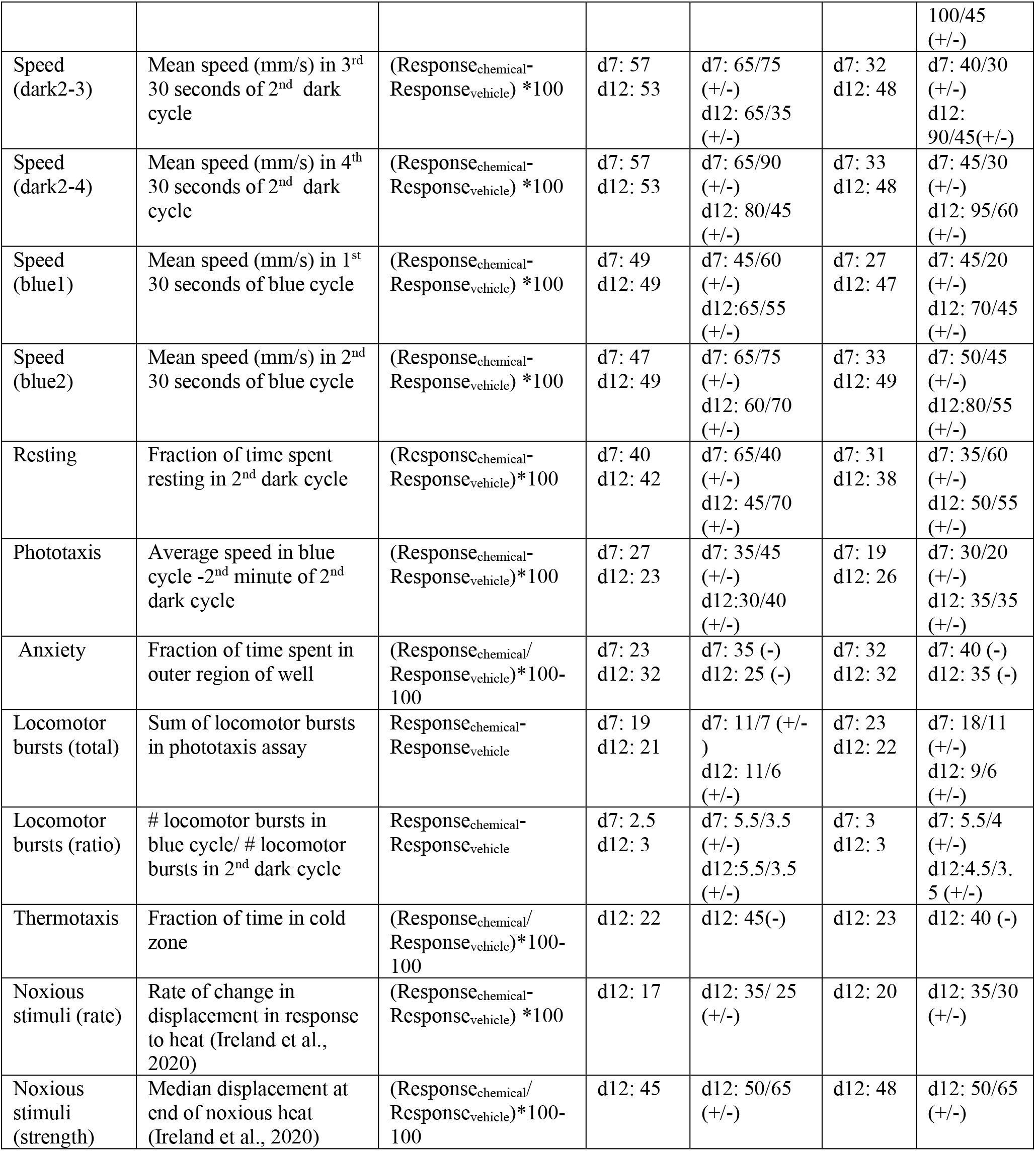
Continuous endpoints. The standard deviation (SD) of the normalized response in the vehicle controls and benchmark response (BMR) are compared for each endpoint on day 7 (d7) and day 12 (d12). Some endpoints can have effects in both the positive and negative directions. In these cases, the BMRs for each direction (increasing (+) or decreasing (-)) are shown.

| Endpoint | Description | Normalization | Adult |  | Regenerating |  |
| --- | --- | --- | --- | --- | --- | --- |
|  |  |  | SD | BMR | SD | BMR |
| Speed (dark1-1) | Mean speed (mm/s) in 1<sup>st</sup> 30 seconds of 1<sup>st</sup> dark cycle | $(\text{Response}_{\text{chemical}} - \text{Response}_{\text{vehicle}}) * 100$ | d7: 58<br>d12: 51 | d7: 90/35 (+/-)<br>d12: 85/45 (+/-) | d7: 37<br>d12: 51 | d7: 60/35 (+/-)<br>d12: 90/60 (+/-) |
| Speed (dark1-2) | Mean speed (mm/s) in 2nd 30 seconds of 1<sup>st</sup> dark cycle | $(\text{Response}_{\text{chemical}} - \text{Response}_{\text{vehicle}}) * 100$ | d7: 62<br>d12: 54 | d7: 110/90 (+/-)<br>d12: 90/45 (+/-) | d7: 37<br>d12: 54 | d7: 60/40 (+/-)<br>d12: 100/60 |
| Speed (green1) | Mean speed (mm/s) in 1<sup>st</sup> first 30 seconds of green cycle | $(\text{Response}_{\text{chemical}} - \text{Response}_{\text{vehicle}}) * 100$ | d7: 60<br>d12: 49 | d7: 70/65 (+/-)<br>d12: 85/45 (+/-) | d7: 34<br>d12: 51 | d7: 50/30 (+/-)<br>d12: 85/60 (+/-) |
| Speed (green2) | Mean speed (mm/s) in 2<sup>nd</sup> 30 seconds of green cycle | $(\text{Response}_{\text{chemical}} - \text{Response}_{\text{vehicle}}) * 100$ | d7: 64<br>d12: 57 | d7: 60/95 (+/-)<br>d12: 95/60 (+/-) | d7: 37<br>d12: 55 | d7: 60/115 (+/-)<br>d12: 115/65 |
| Speed (dark2-1) | Mean speed (mm/s) in 1<sup>st</sup> 30 seconds of 2nd dark cycle | $(\text{Response}_{\text{chemical}} - \text{Response}_{\text{vehicle}}) * 100$ | d7: 52<br>d12: 47 | d7: 50/70 (+/-)<br>d12: 60/30 (+/-) | d7: 26<br>d12: 46 | d7: 35/30 (+/-)<br>d12: 55/50 (+/-) |
| Speed (dark2-2) | Mean speed (mm/s) in 2<sup>nd</sup> 30 seconds of 2<sup>nd</sup> dark cycle | $(\text{Response}_{\text{chemical}} - \text{Response}_{\text{vehicle}}) * 100$ | d7: 57<br>d12: 53 | d7: 60/90 (+/-)<br>d12: 50/55 (+/-) | d7: 29<br>d12: 48 | d7: 40/30 (+/-)<br>d12: |
| Comparative OP neurotoxicity in planarians |  |  |  |  |  |  |
|  |  |  |  |  |  | 100/45<br>(+/-) |
| Speed<br>(dark2-3) | Mean speed (mm/s) in 3<sup>rd</sup><br>30 seconds of 2<sup>nd</sup> dark<br>cycle | (Response<sub>chemical</sub>-<br>Response<sub>vehicle</sub>)*100 | d7: 57<br>d12: 53 | d7: 65/75<br>(+/-)<br>d12: 65/35<br>(+/-) | d7: 32<br>d12: 48 | d7: 40/30<br>(+/-)<br>d12:<br>90/45(+/-) |
| Speed<br>(dark2-4) | Mean speed (mm/s) in 4<sup>th</sup><br>30 seconds of 2<sup>nd</sup> dark<br>cycle | (Response<sub>chemical</sub>-<br>Response<sub>vehicle</sub>)*100 | d7: 57<br>d12: 53 | d7: 65/90<br>(+/-)<br>d12: 80/45<br>(+/-) | d7: 33<br>d12: 48 | d7: 45/30<br>(+/-)<br>d12: 95/60<br>(+/-) |
| Speed<br>(blue1) | Mean speed (mm/s) in 1<sup>st</sup><br>30 seconds of blue cycle | (Response<sub>chemical</sub>-<br>Response<sub>vehicle</sub>)*100 | d7: 49<br>d12: 49 | d7: 45/60<br>(+/-)<br>d12:65/55<br>(+/-) | d7: 27<br>d12: 47 | d7: 45/20<br>(+/-)<br>d12: 70/45<br>(+/-) |
| Speed<br>(blue2) | Mean speed (mm/s) in 2<sup>nd</sup><br>30 seconds of blue cycle | (Response<sub>chemical</sub>-<br>Response<sub>vehicle</sub>)*100 | d7: 47<br>d12: 49 | d7: 65/75<br>(+/-)<br>d12: 60/70<br>(+/-) | d7: 33<br>d12: 49 | d7: 50/45<br>(+/-)<br>d12:80/55<br>(+/-) |
| Resting | Fraction of time spent<br>resting in 2<sup>nd</sup> dark cycle | (Response<sub>chemical</sub>-<br>Response<sub>vehicle</sub>)*100 | d7: 40<br>d12: 42 | d7: 65/40<br>(+/-)<br>d12: 45/70<br>(+/-) | d7: 31<br>d12: 38 | d7: 35/60<br>(+/-)<br>d12: 50/55<br>(+/-) |
| Phototaxis | Average speed in blue<br>cycle -2<sup>nd</sup> minute of 2<sup>nd</sup><br>dark cycle | (Response<sub>chemical</sub>-<br>Response<sub>vehicle</sub>)*100 | d7: 27<br>d12: 23 | d7: 35/45<br>(+/-)<br>d12:30/40<br>(+/-) | d7: 19<br>d12: 26 | d7: 30/20<br>(+/-)<br>d12: 35/35<br>(+/-) |
| Anxiety | Fraction of time spent in<br>outer region of well | (Response<sub>chemical</sub>/<br>Response<sub>vehicle</sub>)*100-100 | d7: 23<br>d12: 32 | d7: 35 (-)<br>d12: 25 (-) | d7: 32<br>d12: 32 | d7: 40 (-)<br>d12: 35 (-) |
| Locomotor<br>bursts (total) | Sum of locomotor bursts<br>in phototaxis assay | Response<sub>chemical</sub>-<br>Response<sub>vehicle</sub> | d7: 19<br>d12: 21 | d7: 11/7 (+/-<br>)<br>d12: 11/6<br>(+/-) | d7: 23<br>d12: 22 | d7: 18/11<br>(+/-)<br>d12: 9/6<br>(+/-) |
| Locomotor<br>bursts (ratio) | # locomotor bursts in<br>blue cycle/ # locomotor<br>bursts in 2<sup>nd</sup> dark cycle | Response<sub>chemical</sub>-<br>Response<sub>vehicle</sub> | d7: 2.5<br>d12: 3 | d7: 5.5/3.5<br>(+/-)<br>d12:5.5/3.5<br>(+/-) | d7: 3<br>d12: 3 | d7: 5.5/4<br>(+/-)<br>d12:4.5/3.<br>5 (+/-) |
| Thermotaxis | Fraction of time in cold<br>zone | (Response<sub>chemical</sub>/<br>Response<sub>vehicle</sub>)*100-100 | d12: 22 | d12: 45(-) | d12: 23 | d12: 40 (-) |
| Noxious<br>stimuli (rate) | Rate of change in<br>displacement in response<br>to heat (Ireland et al.,<br>2020) | (Response<sub>chemical</sub>-<br>Response<sub>vehicle</sub>)*100 | d12: 17 | d12: 35/ 25<br>(+/-) | d12: 20 | d12: 35/30<br>(+/-) |
| Noxious<br>stimuli<br>(strength) | Median displacement at<br>end of noxious heat<br>(Ireland et al., 2020) | (Response<sub>chemical</sub>/<br>Response<sub>vehicle</sub>)*100-100 | d12: 45 | d12: 50/65<br>(+/-) | d12: 48 | d12: 50/65<br>(+/-) |

Histograms of the normalized control responses for all continuous endpoints are shown in Supplementary Figures 3 and 4. As appropriate, the normalized outcome measures were multiplied by 100 to represent the percent change from the control populations and to provide an appropriate range to perform the BMC analysis. This was done for all endpoints, except for the locomotor bursts endpoints, where no scaling factor was used. The normalized data were used as input for the Rcurvep package, which can calculate an appropriate threshold level – benchmark response (BMR) – at which to define a significant deviation from noise levels (Hsieh et al., 2019). For all endpoints except the locomotor burst endpoints, the BMR was calculated by testing thresholds from 5-95 in steps of 5 and by bootstrapping n=100 samples. Because of the smaller response range in the locomotor bursts endpoints, the thresholds were tested in steps of 1 and 0.5 for the total and ratio endpoints, respectively. For endpoints with large variability (resting, speed), the maximum tested threshold was increased to 150 to ensure a stabilized response. We found that using n=100 provided similar results to the recommended n=1000 (Supplementary Figure 5) but generated results in a much shorter amount of time. The recommended BMR was then used to calculate the BMC for each endpoint, which was performed using n=1000 bootstrapped samples. For day 7 lethality in regenerating planarians, the variance was already minimized at the starting test threshold of 5, thus we manually set the BMR to 10. This threshold level corresponds to the level at which significant effects could be detected by a Fisher exact test. The BMR could not be appropriately calculated for stickiness because of issues with control variability and because some chemicals had increased stickiness at all test concentrations. Thus, we used 2.5 SD to manually set the BMR to 50. We also found that for CPF and dichlorvos, partial lethality at the highest test concentration caused a non-monotonic response that disagreed with the rest of the curve. For these chemicals, the highest concentration was masked in the BMC calculation for stickiness. If necessary, data from a replicate run that was inconsistent with the remaining replicates was excluded. For all outcome measures, we report the median BMCs calculated from these bootstrapped results. The lower and upper limits (5^th^ and 95^th^ percentiles, respectively) of the BMC for each endpoint are listed in Supplemental File 1. Some endpoints can be affected in both directions (e.g., increases or decreases in speed). For these endpoints, BMRs and BMCs were calculated for each direction (Table 3).

### 2.6 Hierarchical clustering

To compare the phenotypic signatures of the OPs to those of the mechanistic control compounds, a “phenotypic barcode” was created for each chemical concentration. These barcodes consisted of a binarized score for every outcome measure indicating whether the median compiled score for a concentration was active (beyond the BMR for that outcome measure) or inactive. Note that because this analysis does not consider concentration-response, this binarization is not meant as definitive hit identification on the individual outcome level, but rather provided a means to compare phenotypic patterns. Outcome measures with two possible directions were separated into the positive or negative direction, resulting in 70 and 71 outcome measures for adult and regenerating planarians, respectively. The data were then filtered to only keep chemical concentrations with at least one active outcome measure and outcome measures that had at least one active hit across all chemicals.

Concentrations with 100% lethality were also removed to focus the analysis on sublethal effects. Hierarchical clustering was performed using binary distance and Ward’s method of clustering (ward.D2) using pheatmap in R (R Core Team, 2016). The chemicals were separated into 6 clusters which appeared to have similar phenotypic patterns within each cluster when manually inspected.

### 2.7 Ellman assays

Thirty-six adult planarians were exposed to either 0.5% DMSO or the respective OPs for 12 days (Table 4). Concentrations were chosen to span the breadth of AChE inhibition, from no inhibition to complete inhibition, when possible.

**Table 4.** OP concentrations tested in Ellman assays.

| Chemical Name | Concentrations tested (μM) |
| --- | --- |
| Acephate | 0.0316, 0.316, 3.16, 31.6, 316 |
| Chlorpyrifos | 0.00316, 0.0316, 0.316, 3.16, 10 |
| Diazinon | 0.0001, 0.001, 0.01, 0.178, 0.316, 3.16 |
| Dichlorvos | 0.0001, 0.001, 0.01, 0.1, 0.316, 3.16 |
| Malathion | 0.0316, 0.316, 3.16, 10, 56.2, 100 |
| Parathion | 0.000316, 0.00316, 0.0316, 0.316, 3.16 |
| Profenofos | 0.000316, 0.00316, 0.0316, 0.316, 1.78 |

Planarians were kept in 12-well plates, with 6 planarians per well and a total volume of 1.2 mL of the test solution to keep the ratio of chemical/planarian consistent with the screening set-up. Any fission events or planarians from wells with death were excluded from the assay. After exposure, the planarians were washed 3X with IO water and then homogenized in 1% TritonX-100 in PBS as described in (Hagstrom et al., 2017b, 2018). An Ellman assay (Ellman et al., 1961) was then performed using an Acetylcholinesterase activity assay kit (Sigma-Aldrich). Absorbance was read at 412 nm every minute for 10 minutes using a VersaMax (Molecular Devices, San Jose, CA) spectrophotometer. AChE activity was calculated as the rate of change of absorbance per minute during the linear portion of the reaction. AChE activity was normalized by protein concentration as determined by a Coomassie (Bradford) protein assay kit (Thermo Scientific, Waltham, MA) and compared to 0.5% DMSO exposed samples (set at 100% activity). Activity measurements were performed with at least three technical replicates per condition and at least 2 independent experiments (biological replicates). The inhibition dose response curves were fit to a log-logistic equation (setting the lower limit to 0, the upper limit to 100, and using the IC_50_ as a parameter) using the drc R package (Ritz et al., 2015). The function ED was then used to obtain the concentration that caused 80% inhibition (IC_80_).

## 3 Results

### 3.1 Exposure to the 7 OPs elicits different types of NT/DNT

Adult and regenerating planarians were exposed for 12 days to 10 concentrations each of the 7 OPs (acephate, chlorpyrifos, diazinon, dichlorvos, malathion, parathion, and profenofos) (Figure 1). To focus on sublethal effects, the highest test concentrations were chosen to be at or just below lethal concentrations based on previous data (Zhang et al., 2019a) or preliminary studies (not shown). No lethality was observed in acephate up to the highest soluble concentration (316 µM), thus this was set as the highest concentration.

**Figure 1.**
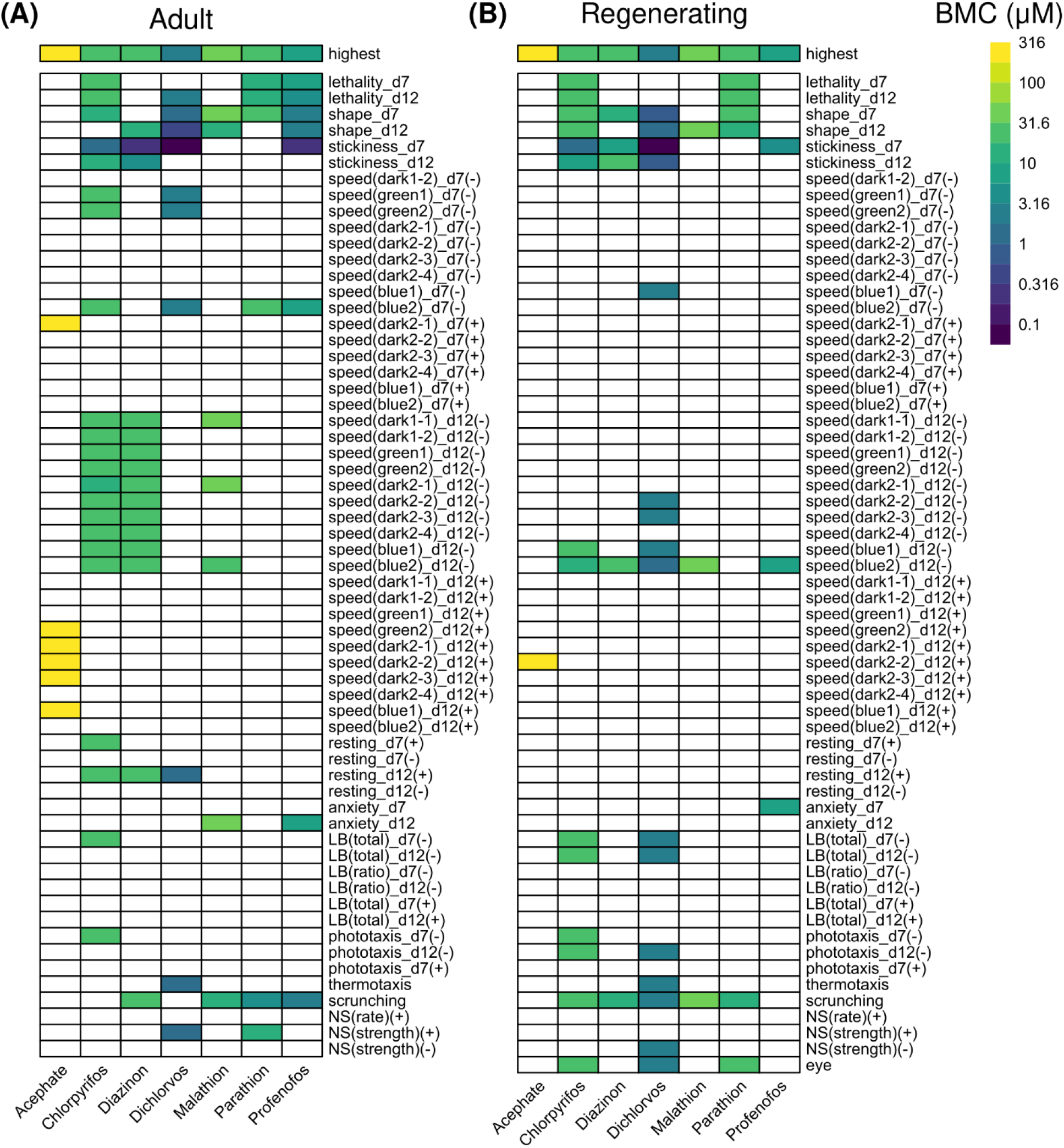
Comparison of OP toxicity. Heatmaps comparing the benchmark concentrations (BMCs) for the OPs in adult (A) and regenerating (B) planarians. The first row shows the highest tested concentration. For outcome measures that can have effects in both directions, the BMCs are separated by either the positive (+) or negative (-) direction. LB: locomotor bursts; NS: noxious stimuli.

The toxicity of the different OPs manifested in different ways (Figure 1). Acephate was the least toxic and caused increased speed (hyperactivity) only at the highest test concentration (316 µM) in adult and regenerating planarians. More speed measures were affected in adults compared to in regenerating planarians. Parathion also caused few hits. In regenerating planarians, effects were only seen at lethal concentrations. Only a defect in scrunching was seen at sublethal concentrations of parathion in adult planarians. In contrast, the remaining 5 OPs showed selective effects in adult and regenerating planarians with morphological and/or behavioral effects in the absence of lethality.

As shown in Figure 2, the endpoints most often affected by OP exposure (at any concentration) were abnormal body shapes, stickiness, scrunching, and speed in the blue light period. These endpoints were largely shared by all the OPs, except for parathion and acephate, in both adult and regenerating planarians. These endpoints were also the most sensitive to detect OP toxicity. Increased stickiness (on day 7) was the most sensitive BMC for 4 OPs (CPF, diazinon, dichlorvos, and profenofos) in both adult and regenerating planarians. Abnormal body shapes on day 12 were the most sensitive BMC for malathion in adult planarians, while decreased speed in the blue light period was the most sensitive endpoint for regenerating planarians. Scrunching was the most sensitive endpoint for adult planarians exposed to parathion.

**Figure 2.**
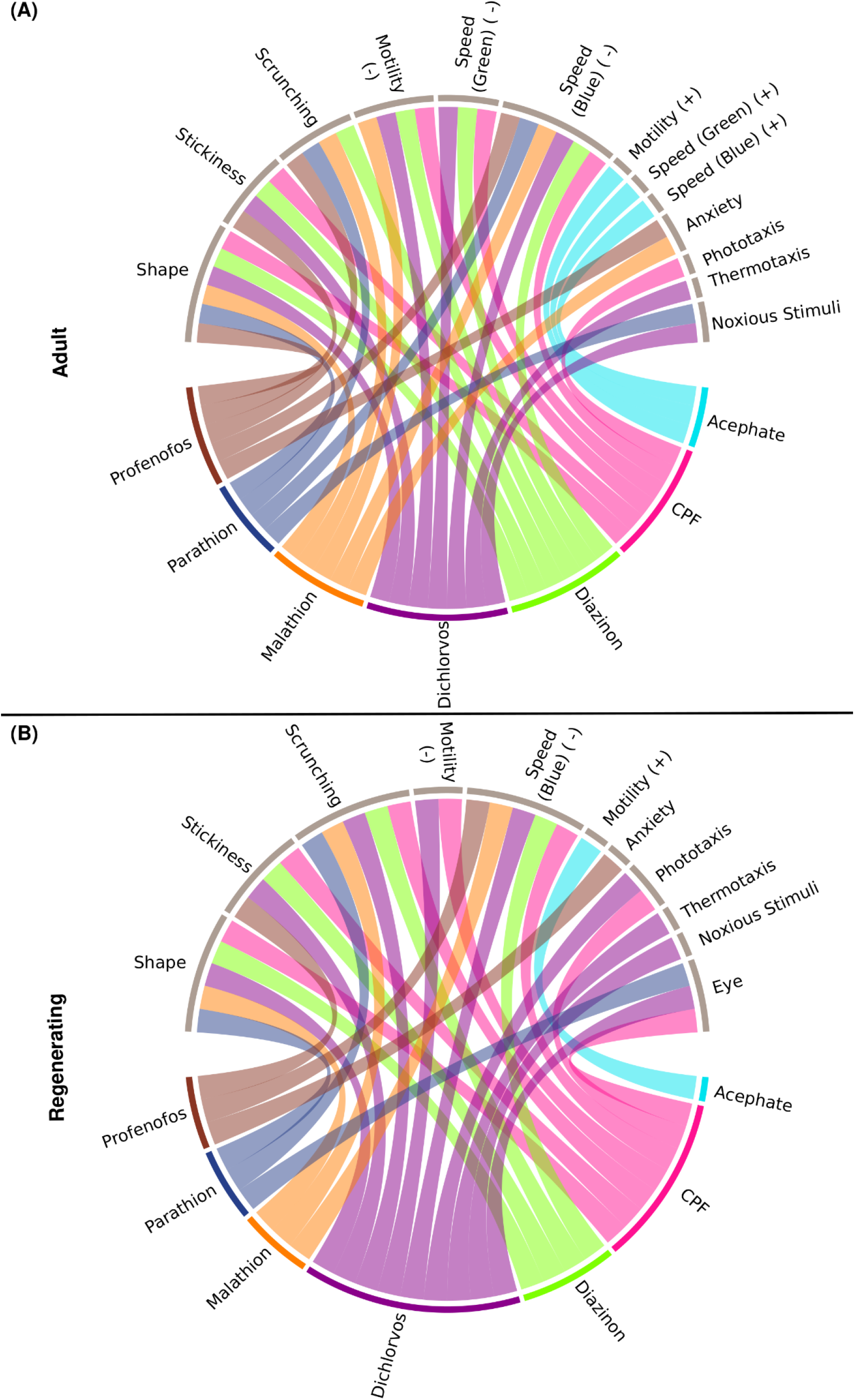
Connections between individual OPs and classes of endpoints. Interaction of the 7 OPs with the different endpoint classes for (A) adult and (B) regenerating planarians. Connections were made if the OP caused a hit at either day 7 or 12 at any tested concentration. Effects on speed in the dark period, resting, or locomotor bursts were combined into the “Motility” category. Speed(B): speed in the blue period, Speed(G): speed in the green period, PT: Phototaxis, NS: noxious stimuli. For endpoints that had effects in both directions, the BMCs are separated by either the positive (+) or negative (-) direction.

Comparing the most sensitive BMC, the potency ranking of the OPs was similar between adult and regenerating planarians (Supplemental Table 1), except that the ranking of diazinon and CPF were swapped in the adult and regenerating planarians. Adults were more sensitive to diazinon than CPF, while regenerating planarians were relatively more sensitive to CPF than diazinon. This change was due to diazinon having a much higher BMC (lower potency) in regenerating planarians compared to adult planarians, since the potency of CPF was the same in the two worm types. Notably, even though no differential overall sensitivity was observed with CPF or dichlorvos between the two worm types, CPF and dichlorvos both affected more/different categories of endpoints in regenerating versus adult planarians (Figure 2). In contrast, adult planarians were more sensitive to diazinon, malathion, and profenofos in terms of overall potency and number of endpoint categories affected compared to regenerating planarians (Supplemental Table 1, Figure 2).

Focusing on sublethal concentrations only, in adult planarians, 5 OPs (CPF, dichlorvos, malathion, profenofos, and diazinon) caused abnormal body shapes, but only 3 of them (diazinon, dichlorvos, and malathion) also caused abnormal body shapes in regenerating planarians. Contraction was the primary body shape associated with OP exposure (Figure 3). In addition, adult and regenerating planarians exposed to dichlorvos exhibited a mixture of contraction, c-shapes and pharynx extrusion on day 7 and contraction and c-shapes on day 12 (Figure 3). Worms that exhibited pharynx extrusion on day 7 were dead by day 12, suggesting that pharynx extrusion was an early indicator of systemic toxicity.

**Figure 3.**
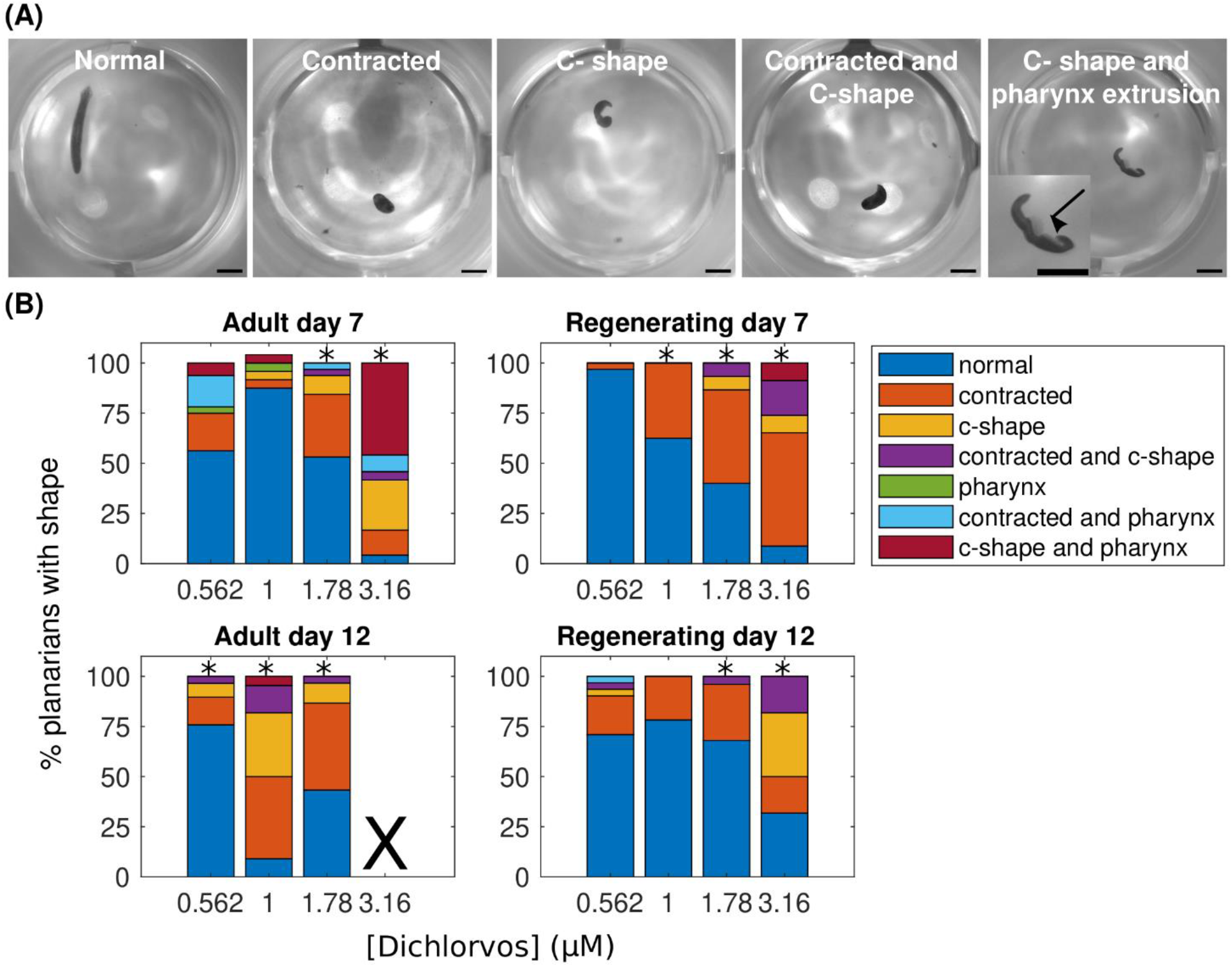
Abnormal body shapes induced by OP exposure. (A) Examples images of normal and abnormal body shapes observed in the OP-treated planarians. The normal planarian is from the vehicle controls. The contracted planarian was treated with 316 µM malathion. The remaining images are of dichlorvos-treated planarians. Scale bar: 1 mm. Inset shows magnified image of the planarian with the arrow pointing to the pharynx which is extruded from the body. (B) Stacked bar plot showing the percentage of planarians exhibiting the different body shape categories as a function of dichlorvos concentration in µM. Concentrations which are above the BMC for the outcome measure are marked with *. X indicates 100% lethality at 3.16 µM dichlorvos in adult planarians on day 12.

### 3.2 AChE inhibition alone cannot explain OP toxicity profiles

Because acetylcholinesterase is a shared target of these OPs, we investigated whether the observed phenotypic differences in adult planarians could be explained by differences in AChE inhibition. Ellman assays were performed on adult planarians exposed for 12 days to different concentrations of the OPs to determine the IC_80_ (Figure 4, Supplementary Figure 6). We calculated the IC_80_ because significant cholinergic toxicity is seen in mammals when AChE inhibition reaches about 80% (Lionetto et al., 2013; Russom et al., 2014; Voorhees et al., 2017). Comparisons were made to the most sensitive BMC in adult planarians (BMC_adult_). Significant inhibition was not observed in up to 316 µM acephate. For malathion, significant lethality was observed before 80% inhibition could be reached and the extrapolated IC_80_ was significantly higher than the lowest BMC_adult_ (Figure 4). For dichlorvos, the IC_80_ was also higher than the most sensitive BMC_adult_. For the remaining 4 OPs, the IC_80_ values were all lower (CPF, profenofos, parathion) or close to (diazinon) the most sensitive BMC_adult_. Diazinon and dichlorvos had very similar inhibition profiles but very different toxicity profiles. Given the differences in observed phenotypic outcomes, this suggests that AChE inhibition alone cannot explain the manifestation of toxic outcomes (Figure 4). Similarly, no correlation was observed between OP toxicity and hydrophobicity (logP, Figure 4A). Acephate is the only hydrophilic OP and neither showed significant behavioral effects nor AChE inhibition.

**Figure 4.**
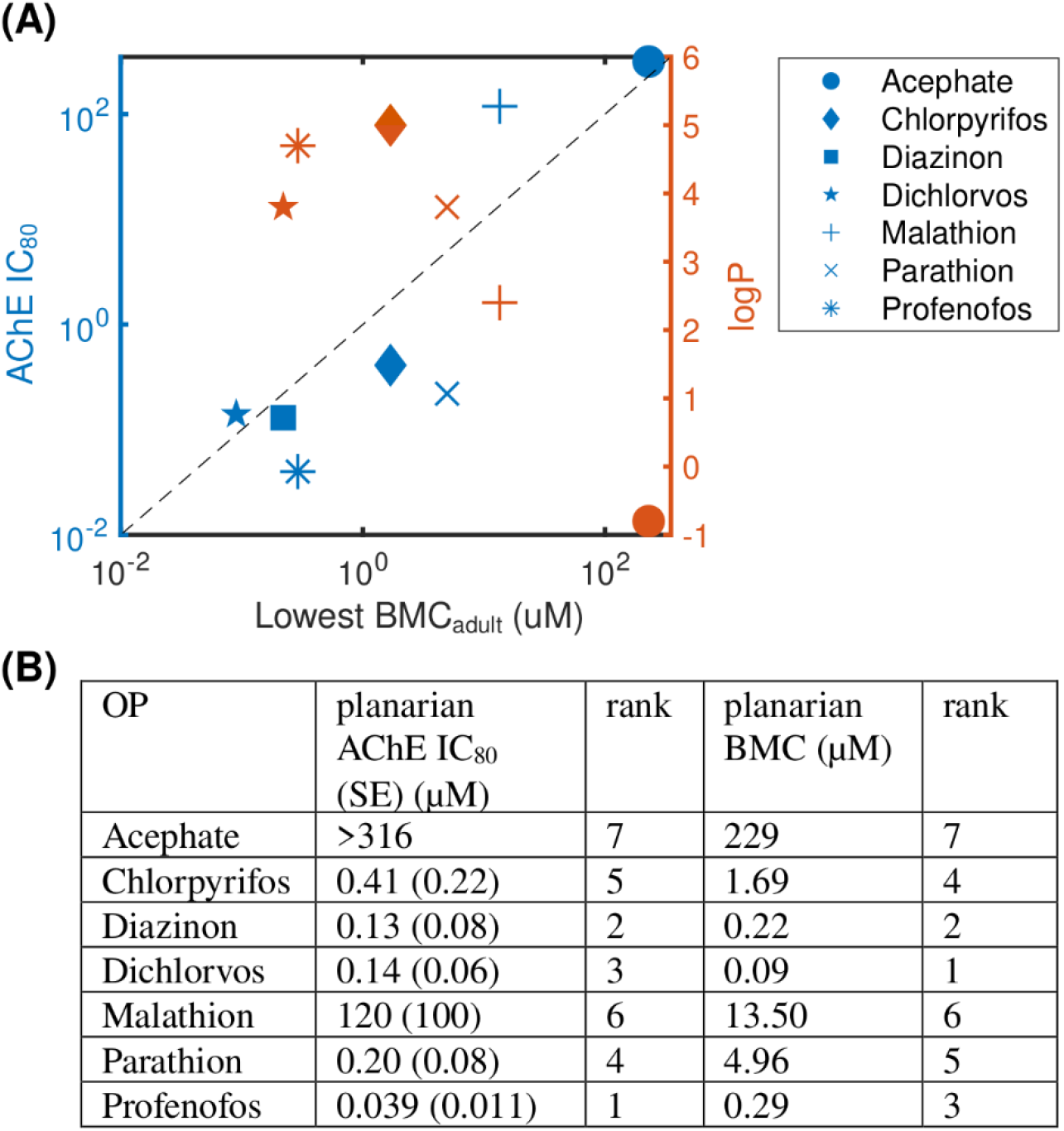
Levels of AChE inhibition is not correlated with phenotypic effects. (A) Comparison of OP potency (measured by the most sensitive BMC_adult_) to AChE inhibition (IC_80_, blue) and hydrophobicity (logP, orange). Raw data and fitted curves from the Ellman assays are shown in Supplementary Figure 6. For acephate, the IC_80_ was set to the highest test concentration since no inhibition was observed. The dashed line is provided as a visual tool to show when the IC_80_ equals the lowest BMC_adult_. Blue points above this line (malathion and dichlorvos) indicate the BMC was more sensitive than the IC_80_. (B) A comparison of the ranking of the potency of the 7 OPs using the planarian AChE IC_80_ and most sensitive BMC_adult_.

To compare results across the OPs, the potency of the 7 OPs was ranked using either the BMC_adult_ or the IC_80_ (Figure 4B). Overall trends were similar when comparing rankings across the two measures (Spearman’s correlation coefficient: 0.82; p-value: 0.03), with diazinon, dichlorvos, and profenofos being the most potent OPs and acephate and malathion the least potent in both measures. Profenofos, however, was the most potent OP using AChE inhibition whereas dichlorvos was the most potent when looking at the most sensitive BMC_adult_. CPF and dichlorvos showed higher relative potency rankings when measured by the most sensitive BMC_adult_ compared to the IC_80._ The potency ranking found here for day 12 adult IC_80_ was similar to ranking comparing inhibition rates of planarian homogenates using the oxons of a subset of the OPs tested here (CPFO, diazoxon, dichlorvos, paraoxon and malaoxon) (Hagstrom et al., 2017a).

### 3.3 Comparison of OP toxicity with known effectors of mechanistic pathways

To delineate whether certain OPs affect other targets besides AChE, we also screened “mechanistic control chemicals” known to target pathways that have been implicated in OP DNT. These chemicals were screened in 5 concentrations each in adult and regenerating planarians and the BMCs were calculated (Supplementary Figure 7).

To determine whether any phenotypic patterns could be discerned in the data, we generated phenotypic barcodes for each chemical concentration consisting of a binary score (active or inactive) for each outcome measure. Hierarchical clustering was performed on the binarized phenotypic barcodes to see whether different chemical concentrations shared phenotypic patterns. We chose to look at individual concentrations because different toxicities may emerge depending on the concentration. In both adult (Figure 5) and regenerating (Figure 6) planarians, we identified 6 main clusters based on their phenotypic patterns. Using the mechanistic control compounds, we then tried to anchor the phenotypic clusters to potential underlying mechanisms.

**Figure 5.**
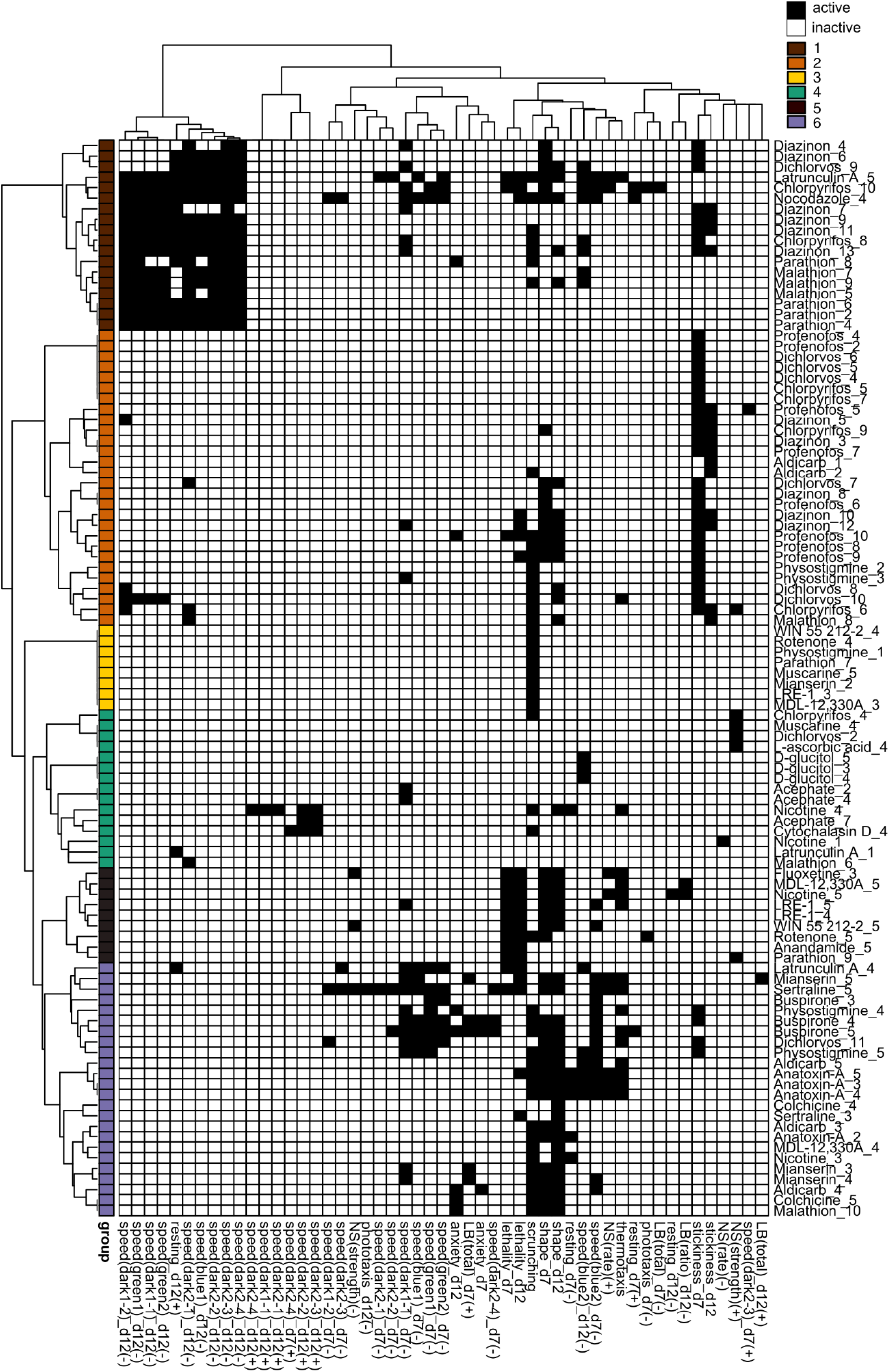
Hierarchical clustering of adult phenotypic barcodes. Active outcome measures for each chemical concentration are shown in black. Outcome measures that can have effects in both directions are separated into either the positive (+) or negative (-) direction. Only active chemical concentrations and outcome measures are shown. Numbers in chemical names refer to the respective test concentration, with 1 as the lowest tested concentration, see Table 1. Hierarchical clustering was performed using binary distance and Ward’s method. Six clusters were identified with similar phenotypic profiles: Cluster 1: Strong locomotor defects (reduced speed and increased resting), Cluster 2: effects in primarily day 7 stickiness, Cluster 3: scrunching defects only, Cluster 4: 1 or 2 hits in miscellaneous outcome measures, Cluster 5: lethal/systemic toxicity, Cluster 6: effects in scrunching and body shape with the addition of other outcomes.

**Figure 6.**
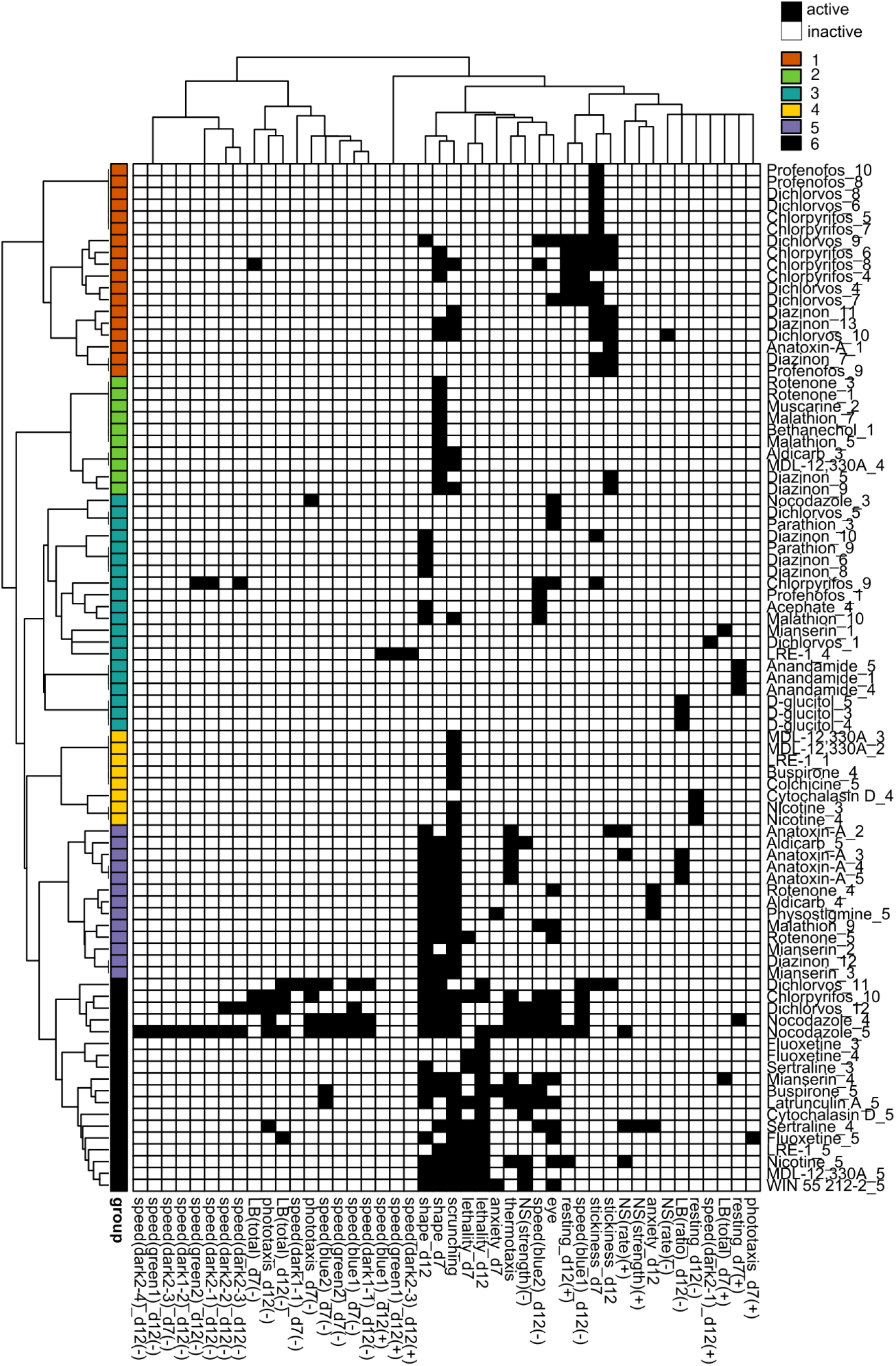
Hierarchical clustering of regenerating phenotypic barcodes. Active outcome measures for each chemical concentration are shown in black. Outcome measures that can have effects in both directions are separated into either the positive (+) or negative (-) direction. Only active chemical concentrations and outcome measures are shown. Numbers in chemical names refer to the respective test concentration, with 1 as the lowest tested concentration, see Table 1. Hierarchical clustering was performed using binary distance and Ward’s method. Six distinct clusters were identified with similar phenotypic profiles: Cluster 1: effects on stickiness, Cluster 2: abnormal body shapes, Cluster 3: hits in miscellaneous endpoints, Cluster 4: primarily scrunching defects, Cluster 5: effects in scrunching and body shape with the addition of other outcomes, Cluster 6: lethal/systemic toxicity.

In adult planarians (Figure 5), several mid-to high range concentrations of diazinon, malathion, parathion, and CPF were found in cluster 1, indicating locomotion defects. Only high concentrations of latrunculin A and nocodazole were also found in this cluster. Low to mid-range concentrations of CPF, diazinon, dichlorvos, malathion, and profenofos were found in cluster 2, which was characterized by effects in day 7 stickiness. The carbamate AChE inhibitors physostigmine and aldicarb as well as the nicotinic AChR agonist nicotine were also associated with this cluster. Single concentrations of various chemicals, including a concentration of parathion, were found in cluster 3, with specific effects in scrunching only. Cluster 4 affected miscellaneous outcome measures and contained concentrations of the negative control D-glucitol, three concentrations of acephate, and single random concentrations of CPF, muscarine, malathion, and dichlorvos. Small subclusters appear to emerge within this cluster. For example, one of the concentrations of acephate (7) is associated with hyperactive effects (increased speed) along with one concentration each of nicotine and cytochalasin D. High concentrations of dichlorvos and malathion were associated with cluster 5, which was characterized by effects on scrunching and body shape, often with the addition of various other outcome measures. This cluster also contained the higher concentrations of several cholinergic compounds (aldicarb, physostigmine, anatoxin-A and nicotine), serotonergic compounds (buspirone, mianserin, sertraline) and cytoskeletal disrupting compounds (colchicine, latrunculin A), suggesting this collection of phenotypes may represent convergence of several pathways. Cluster 6 was indicative of lethality and contained high concentrations of various compounds, including OPs.

We observed different clusters of the OPs in regenerating planarians (Figure 6) than in adults (Figure 5). In regenerating planarians (Figure 6), low-to mid-range concentrations of CPF, diazinon, dichlorvos and profenofos were associated with cluster 1, which also contained low concentrations of anatoxin-A and was characterized by increased stickiness. Effects on day 12 speed(blue1) and day 12 resting were also associated with this cluster. Mid-range concentrations of diazinon and malathion were associated with cluster 2, which was characterized by abnormal body shapes on day 7. Low-range concentrations of dichlorvos and profenofos and higher concentrations of CPF, and diazinon were associated with cluster 3 and displayed miscellaneous hits in a handful of various outcome measures. A single concentration of acephate was also found in this cluster. High concentrations of malathion and diazinon were associated with cluster 5, which was characterized by effects on body shape and scrunching. This cluster also contained the higher concentrations of several cholinergic compounds (aldicarb, physostigmine, anatoxin-A), the serotonin receptor agonist mianserin, and the oxidative stress inducer rotenone. Lastly, cluster 6 was associated with lethality and contained the highest concentrations of CPF and dichlorvos. None of the OPs were associated with cluster 4.

## 4 Discussion

### 4.1 OPs cause distinct toxicity profiles in adult and regenerating planarians

Previous studies have used freshwater planarians to study the effects of OP exposure on survival, behavior, and regeneration (Levy and Miller, 1978; Villar et al., 1993; Feldhaus et al., 1998; Zhang et al., 2013; Poirier et al., 2017; Hagstrom et al., 2018). However, these studies have focused only on a few endpoints and/or compounds and have been largely qualitative. This study investigated the effect of 7 OPs on adult and regenerating planarians in parallel using multiple quantitative phenotypic readouts to assay sublethal effects of continuous OP exposure. We found that these 7 OPs had distinct toxicity profiles in planarians that differed across the OPs and between adult and regenerating individuals, recapitulating findings in mammalian systems showing that OPs cause a variety of toxic outcomes (reviewed in (Moser, 1995; Pope, 1999; Voorhees et al., 2017)).

In terms of potency, regenerating planarians were overall less sensitive to the OPs. At the tested concentrations, lethality was only observed for 2 OPs (CPF and parathion) in regenerating planarians, whereas 4 OPs (CPF, parathion, dichlorvos and profenofos) showed significant lethality in adult planarians by day 12. This increased lethality in adult versus regenerating planarians was also observed in the mechanistic control compounds tested here and has been observed for other chemicals (Hagstrom et al., 2015; Zhang et al., 2019a). Metabolic differences in the two developmental stages may explain some of the differential sensitivity to chemical exposure. Oxygen consumption and metabolism are slower in larger planarians than in smaller ones (Hyman, 1919; Osuma et al., 2018). To ensure similar sizes after amputation of the regenerating group to the adult group, we amputate larger planarians to generate the regenerating group, thus regenerating planarians may have slower metabolism. In addition, while it was found that oxygen consumption is not different between resting intact/adult and regenerating planarians of similar size (Osuma et al., 2018), oxygen consumption depends on activity levels (Jenkins, 1959) and regenerating planarians are less mobile during the first 5-7 days after exposure/amputation than adults (Zhang et al., 2019a). It is also possible that the pharmacokinetics differ between regenerating and adult planarians, resulting in potentially lower internal doses in regenerating planarians. These differences could be chemical-specific. Future work characterizing the pharmacokinetics in both developmental stages will be critical to contextualize results in planarians and for dose comparisons with other species.

To prepare for this kind of system comparison of our results, we used BMC analysis, in contrast to earlier studies that use lowest-observed-effect levels (LOELs), and thus are dependent on the exact test concentrations. Typically, point-of-departure/BMC analysis has relied on the use of fixed threshold levels, e.g., 2 or 3 SD, to determine effects. This approach is not always appropriate for behavioral data that is often not normally distributed. Thus, we utilized the Rcurvep package which determines the appropriate threshold/BMR based on the actual noise in the data, which had previously been used with developing zebrafish morphological and behavioral data (Hsieh et al., 2019). A comparison of SD and BMR values is provided in Tables 2 and 3 and shows that for most endpoints the BMR is around 2-3 SDs. Of note, this approach assumes a monotonic concentration response, and thus is not as sensitive to non-monotonic effects, potentially decreasing sensitivity for some chemicals compared to our previous studies using LOELSs (Zhang et al., 2019a). The hierarchical clustering in Figures 5 and 6 does not take into consideration concentration-response; therefore, non-monotonic hits are observed in these phenotypic profiles that were not detected by the BMC analysis and thus their biological significance is unclear. Determination of the BMRs revealed a surprising level of phenotypic diversity within the morphologies and behaviors of both control and chemically-treated planarians evidenced by a large SD/BMR of some endpoints. This variability suggests that although asexual planarians are considered clonal, there is differential sensitivity to physical and chemical stimuli. Similar variability in the behavioral response of individual *D. japonica* planarians has previously been reported when planarians were simultaneously exposed to two stimuli (a chemoattractant and bright light) (Inoue et al., 2015). This phenotypic diversity may reflect the genetic diversity that has been observed in genes involved in responses to external stimuli, such as genes associated with “signal transduction” and “defense mechanisms” in asexual *D. japonica* planarians (Nishimura et al., 2015). In contrast to highly in-bred animal models, this genetic diversity may be beneficial to more accurately reflect the large amount of variability found in the diverse human population, where the goal is to protect the most sensitive populations.

### 4.2 AChE inhibition is not predictive of OP toxicity in adult planarians

Comparison of the presence of adverse outcomes (most sensitive BMC_adult_) with AChE inhibition (IC_80_) demonstrated that significant AChE inhibition on day 12 was not predictive of phenotypic effects in adult planarians (Figure 4). The OPs showed different relative ranking when comparing IC_80_s to the most sensitive BMC_adult_. While the IC_80_ for profenofos was ∼3X lower than that of dichlorvos, the BMC_adult_ for dichlorvos was 3X lower than that of profenofos. Similarly, the IC_80_ for parathion was half of that for CPF, but the most sensitive BMC_adult_ for CPF was about 4 times lower than that of parathion. Dichlorvos, malathion and acephate showed phenotypic effects at concentrations below their IC_80._ Diazinon had phenotypic effects emerge at approximately the IC_80_, whereas effects were only seen above the IC_80_ for CPF, parathion, and profenofos. For the latter group, the IC_80_s were 4x (CPF) to 23x (profenofos) lower than the respective BMCs. This means that although AChE is significantly inhibited at low concentrations, morphological or behavioral effects were not observed.

In humans, 50-60% AChE inhibition produces mild symptoms, 60-90% inhibition produces moderate symptoms, and >90% inhibition causes death due to respiratory or heart failure (Lionetto et al., 2013). Since planarians lack target organs such as lungs and a heart, it is possible that even >90% inhibition does not cause death as rapidly as in humans. Similar trends showing no correlation between AChE inhibition and lethality has also been shown in nematodes (Rajini et al., 2008).

However, some phenotypic effects would still be expected at these high levels of inhibition. The fact that we were able to achieve complete loss of AChE activity (within the detection limit of our assay) with CPF, parathion and profenofos in the absence of phenotypic effects suggests that 1) in the planarian system the enzymatic activity of AChE is not a good biomarker for measuring the adverse effects of OP exposure, 2) that some effects due to high AChE inhibition only manifest after the day 12 screening, or 3) that planarians have compensatory mechanisms that allow them to adjust to low dose OP exposure over time. In support of this last idea, we found that stickiness, which is a sensitive endpoint for cholinergic effects (Hagstrom et al., 2018), was a shared and sensitive endpoint affected by many of the OPs on day 7. However, stickiness on day 12 was not affected to the same extent and often had lower sensitivity. We have previously shown that short term (5 day) exposure to physostigmine and diazinon caused increased stickiness which was not seen in day 12 regenerating planarians, despite nearly complete inhibition of AChE activity in both worm types (Hagstrom et al., 2018). In support of this idea that compensatory mechanisms may protect from cholinergic toxicity, it has been shown that compensatory down-regulation of muscarinic and nicotinic ACh receptors occurs with many OPs, including parathion. This can lead to tolerance of long-term inhibition of AChE, but tolerance can vary with different OPs (Bushnell et al., 1993; Jett et al., 1993; Dvergsten and Meeker, 1994; Voorhees et al., 2017; Slotkin et al., 2019). Similarly, compensatory down-regulation of nicotinic AChRs and morphological remodeling have been shown to occur in the AChE -/- knockout mouse, which can surprisingly survive until adulthood despite complete loss of AChE activity (Chatonnet et al., 2003; Adler et al., 2004). Future mechanistic studies will need to evaluate regulation of receptor gene expression to address these possibilities and link phenotypes to molecular events.

### 4.3 Clustering with mechanistic control chemicals contextualizes OP toxicity profiles

Comparison of the phenotypic profiles of the different OPs with those of the mechanistic control compounds was used to parse out effects on cholinergic and non-cholinergic targets. As expected, some of the phenotypic outcomes were shared by many of the OPs, reflecting their shared action on AChE. Four OPs (profenofos, CPF, diazinon, and dichlorvos) caused increased stickiness at day 7, which were also found with exposure to the carbamates aldicarb and physostigmine (Figure 5 cluster 2, Figure 6 cluster 1). None of the other control compounds caused increased stickiness, together suggesting increased stickiness may be a sensitive, specific cholinergic effect. We have shown that increased stickiness is correlated with increased mucus secretion (Malinowski et al., 2017). Increased secretions (including bronchial, lacrimal, salivary, sweat, and intestinal secretions) are a major hallmark of acute cholinergic toxicity due to stimulation of muscarinic AChRs (Pope et al., 2005b; Peter et al., 2014; Taylor, 2018).

Defects in scrunching – a musculature-driven gait (Cochet-Escartin et al., 2015) - may also be an indicator of cholinergic toxicity, possibly through effects on nicotinic receptors which are known to regulate motor functions. We previously found that scrunching defects were correlated with AChE inhibition in organophosphorus flame retardants (Zhang et al., 2019b). In the current screen, we observed the inclusion of scrunching in clusters 2 and 6 (adult) and 5 and 6 (regenerating) which contained medium to high concentrations of dichlorvos, profenofos, and malathion in adults and diazinon, CPF, and malathion in regenerating planarians, as well as compounds such as aldicarb, nicotine, physostigmine and anatoxin-A. Notably, these clusters also contained chemicals targeting the serotonergic pathway (mianserin, sertraline, fluoxetine). Modulation of cholinergic signaling by serotonin has been supported by several studies (Cassel and Jeltsch, 1995), and CPF and diazinon have been shown to directly target the serotonin receptors in rodents (Slotkin et al., 2006b, 2019). Uncovering whether and how these OPs modulate serotonergic signaling in planarians would be an interesting next step to investigate.

Scrunching involves many steps from the initial noxious heat sensation to the stereotypical muscle-driven periodic body length oscillations. While our previous work has begun to delineate the sensing mechanisms controlling scrunching in response to diverse physical (e.g., amputation, ultraviolet light) and chemical stimuli (e.g., allyl isothiocyanate, 1% ethanol) (Cochet-Escartin et al., 2015, 2016; Sabry et al., 2019, 2020), we do not yet know what controls scrunching in response to noxious heat. Given that scrunching (in contrast to the default planarian gliding gait which is ciliary driven (Rompolas et al., 2013)) requires coordinated muscle movement (Cochet-Escartin et al., 2015), it is conceivable that it depends on cholinergic signaling. Additionally, we have shown that susceptibility to heat is affected in short-term exposure to diazinon and physostigmine as well as when planarian cholinesterase is knocked down via RNA interference (Hagstrom et al., 2018). Thus, heat sensation could also be affected, with or without impairment of the scrunching motion. Adult AChE -/- knockout mice exhibit decreased pain perception (Duysen et al., 2002). Transient receptor potential (TRP) channels are important for sensing many noxious and painful stimuli and we have shown that several TRP channels are important for inducing scrunching in response to specific stimuli (Sabry et al., 2019). Thus, we speculate that effects on noxious heat sensation in planarians may be a result of downstream effects on TRP channels, though which specific family are responsible remains to be determined. To delineate whether OPs cause defects in heat sensation and/or in scrunching execution, low-throughput mechanistic studies using other scrunching inducers, such as amputation (Cochet-Escartin et al., 2015) will be necessary. Given the complexity of the scrunching pathway, it is expected that not all scrunching defects are specific to cholinergic toxicity. Indeed, cluster 3 (adults) and cluster 4 (regenerating) included various compounds that only affected scrunching (with the exception of nicotine in regenerating planarians in cluster 4), indicating that multiple pathways converge to cause scrunching defects.

CPF, diazinon, dichlorvos, malathion and profenofos all induced abnormal body shapes in adult planarians at sublethal concentrations. Except for CPF and profenofos, these OPs also induced sublethal abnormal body shapes in regenerating planarians. The primary body shape observed was contraction, with dichlorvos also exhibiting a mix of contraction and c-shapes. The nicotinic AChR agonist anatoxin-A also induced a severe contracted phenotype, while a mixture of contracted and c-shape phenotypes was observed in adult and regenerating planarians exposed to aldicarb, physostigmine and nicotine (Supplementary Figure 8). Previous studies have observed increased C-shape hyperkinesia in planarians after acute exposure to nicotine (Rawls et al., 2011). Contraction and our joint classification of contracted/c-shape are equivalent to the “walnut” and “bridge-like” body shapes that have previously been shown to be induced by nicotine and physostigmine, respectively, in the planarian *Dugesia gonocephala* (Buttarelli et al., 2000). In *D. japonica,* physostigmine-induced contraction was shown to be a result of contraction of the body-wall muscles and that these contractions were delayed by preincubation with the muscle nicotinic AChR antagonist tubocurarine or the muscarinic AChR antagonist atropine (Nishimura et al., 2010). Together, these findings suggest that these contracted body shapes may be indicative of cholinergic toxicity.

Looking at the OPs individually in more detail, we observed that profenofos caused few additional effects at sublethal concentrations, suggesting that cholinergic toxicity is the most sensitive effect for profenofos in planarians, similar to findings in other systems (Levy and Perron, 2016). In contrast, CPF, dichlorvos and diazinon affected multiple different endpoints in adult and regenerating planarians. We observed phenotypic effects of CPF only at concentrations at which planarian AChE was also significantly inhibited, which may suggest that AChE inhibition drives the toxicity profile of CPF in planarians. However, while adult and regenerating worms had the same lowest BMC, they exhibited different phenotypic profiles, implying that CPF must affect additional different targets in the two developmental stages concurrently with significant AChE inhibition. Similarly, in developing zebrafish, it was found that behavioral defects induced by CPFO were caused by issues with axonal outgrowth and neuronal activity, but only at concentrations of significant AChE inhibition (Yang et al., 2011). In a previous screen (Zhang et al., 2019a), we found that the LOEL for CPF was lower in regenerating planarians (10 µM) than in adults (100 µM). In the present screen, we do not see that potency difference when considering all outcome measures collectively, as log_10_(BMC_adult_/BMC_regenerating_) is zero. However, the previous screen did not evaluate stickiness, which we found to be the most sensitive endpoint for CPF in both worm types with a BMC of 1.7 µM. Furthermore, when looking at an individual outcome-level, developmental selectivity was observed, because certain outcomes (e.g., scrunching) were only affected in regenerating but not adult planarians, consistent with previous results (Zhang et al., 2019a).

A similar trend was observed for dichlorvos, where overall sensitivity was the same in the two developmental stages, but a greater breadth of endpoints was affected in regenerating planarians. Dichlorvos was the most potent OP in adult and regenerating planarians. In adults, phenotypic effects were present at concentrations below the IC_80_. Dichlorvos does not require bioactivation to inhibit AChE and thus may be fast acting. Nematodes have also been shown to be very sensitive to dichlorvos (Cole et al., 2004) and dichlorvos is a useful anthelminthic (Chavarría et al., 1969), suggesting that worms may be especially sensitive to this OP. Dichlorvos was the only OP that induced a multitude of different body shapes at higher concentrations, including contraction, c-shapes, and pharynx extrusion. Pharmacological studies in planarians have suggested that the disruption of specific neurotransmitter systems leads to stereotypical body shapes. For example, drugs that activate cholinergic signalling induce fixed postures such as contraction while drugs that activate dopamine signalling produce screw-like hyperkinesia (Buttarelli et al., 2000, 2008). Thus, compounds that can target multiple neurotransmitter systems may show mixed phenotypes.

Additionally, the manifestation of different body shapes can progress with increasing concentrations. For example, at lower concentrations dichlorvos primarily induced contractions, similar to the other OPs, but at higher concentrations, the body shapes also consisted of c-shapes, mixed contraction/c-shapes akin to “bridge-like hypokinesia” (Buttarelli et al., 2000) and pharynx extrusion. Since c-shapes have also been observed after acute exposure to nicotine (Rawls et al., 2011), to D2 dopamine receptors agonists (Venturini et al., 1989), or to the biocide methylisothiazolinone (Van Huizen et al., 2017), it is unclear whether this progression of shapes represents increased cholinergic toxicity or potentially effects on different targets that only manifest at higher concentrations. Unlike the distinct postures like contraction and c-shapes which seem to represent disrupted neurotransmission and are often reversible (Buttarelli et al., 2008), pharynx extrusion is a morphological change that may be an early indicator of systemic toxicity. In support of this idea, planarians that exhibited pharynx extrusion on day 7 were dead by day 12. Thus, classification of abnormal planarian body shapes and morphologies provides insight into the types of toxicity observed, but further work is needed to connect effects on specific neurotransmitter systems to specific morphologies.

In contrast to CPF and dichlorvos, adult planarians were much more sensitive to diazinon than regenerating planarians, as the lowest BMC_adult_ was 42X lower than the lowest BMC_regenerating_. This difference was largely driven by effects on day 7 stickiness in adults, which was over an order of magnitude lower than the next most sensitive outcome (0.2 µM compared to 4.8 µM). Moreover, in adult worms, diazinon was associated with cholinergic toxicity and extreme locomotor defects, which may be indicative of systemic toxicity. Systemic toxicity was not found in diazinon-exposed regenerating planarians at the tested concentrations. While the effects in adult planarians were similar in CPF-exposed and diazinon-exposed planarians, regenerating planarians were less sensitive to diazinon than to CPF, both in terms of overall potency and in the number of outcomes affected. Regenerating planarians exposed to diazinon primarily only showed effects on the cholinergic-related endpoints (stickiness, body shape, and scrunching). It will be interesting to delineate in future work why regenerating planarians are more resistant to diazinon than adults.

For malathion, effects on body shape and scrunching were among the most sensitive endpoints. Malathion was unable to cause >75% inhibition of planarian AChE without lethality, suggesting that malathion may affect alternative targets more strongly than AChE in planarians. Unlike the previous OPs, we also did not observe any effects on stickiness with malathion. In embryos of *Xenopus laevis* frogs, malathion has been shown to bind with high affinity to lysyl oxidase, causing post-translational modification of collagen and developmental defects (Snawder and Chambers, 1993). Malathion has also been reported to inhibit lysyl hydroxylase in rat cell culture (Samimi and Last, 2001) and cause oxidative damage in rat brain regions (Fortunato et al., 2006). Absence of significant AChE inhibition and cholinergic toxicity when exposed to chronic low doses of malathion has also been reported in rodents and behavioral effects (e.g., memory impairment) have been attributed to alternative effects, such as mitochondrial dysfunction (Trevisan et al., 2008; dos Santos et al., 2016). In regenerating planarians, malathion was associated with cluster 5 which also contained two concentrations of rotenone, a mitochondrial disruptor which leads to increased oxidative stress. Both compounds induced abnormal body shapes and disrupted scrunching.

The remaining two OPs – acephate and parathion - caused strikingly different toxic outcomes. Acephate, a phosphoramide, was the least toxic of all the OPs tested. We only observed hyperactive behavior at the highest test concentration. Acephate is highly water soluble (negative logP; Figure 4) and thus may not easily pass through the planarian epithelium, potentially explaining why it caused few effects. Interestingly, acephate has also been reported to cause hyperactivity in developing zebrafish at 0.1 mg/L concentration (Liu et al., 2018), indicating that there may be a conserved mechanism. Hyperactive effects were also observed in adult planarians exposed to 316 µM nicotine, which closely clustered with acephate (Figure 5, cluster 4). Low concentrations of nicotine have previously been shown to induce hyperactivity and seizure-like motion in planarians (Rawls et al., 2011; Pagán et al., 2015), similar to effects observed in rodents (Zhu et al., 2012). Moreover, acephate proved to be a weak planarian AChE inhibitor as no significant AChE inhibition was observed within the solubility limit of acephate. In humans, acephate also shows weak toxicity, requiring high concentrations to achieve significant AChE inhibition (Ando and Wakamatsu, 1982).

In contrast, we found that parathion was a potent planarian AChE inhibitor and caused lethality in adult and regenerating planarians in the absence of phenotypes at sublethal concentrations, except for scrunching in adult planarians. A similar toxicity profile for parathion (lethality before neurotoxic effects could be detected) has been observed in neonatal rats when treated subcutaneously with parathion (Slotkin et al., 2006a).

In summary, this study shows that behavioral HTS in freshwater planarians is a suitable platform to study OP neurotoxicity and DNT. Bioactivation of OPs occurs at all stages of development in planarians and our results recapitulate findings from mammalian studies showing that different OPs cause different toxicity profiles that cannot be explained by AChE inhibition alone. Adverse outcomes differ between adult and developing organisms due to the OPs’ effects on multiple targets. Thus, this work emphasizes the need for more comparative studies across developmental stages to better understand how different OPs – alone or in combination, as in real life applications – damage the nervous system. The screen presented here is a first step in delineating the different neurotoxic outcomes induced by OP exposure in planarians. The phenotypic clusters that we identified can be used as a starting point for future mechanistic studies of OP neurotoxicity in planarians. Using targeted chemical and genetic (RNAi) screening to manipulate the proposed pathways connected to a specific OP or phenotype will allow for determination of whether this pathway mitigates the observed effects. Such an approach has been successfully used to show that activation of D1 but not D2 dopamine receptors are involved in planarian screw-like hyperkinesia, whereas D2 but not D1 receptors are involved in inducing c-shapes (Venturini et al., 1989). Proteomics and/or RNA-seq studies would be useful to complement targeted screens to verify the targets and pathways affected by different OPs in planarians. The unique strengths of planarian HTS – the ability to directly compare the toxic outcomes of adult and developing animals using the same assays and metrics – make this type of mechanistic analysis particularly powerful, as it would allow for direct connections to be made between molecular targets and adverse outcomes across different developmental stages.

## 6 Author Contributions

DI contributed to experimental design, performed screening experiments, Ellman assays, data analysis and interpretation, and graphical representation. SQ co-maintained the screening platform, performed screening experiments and data analysis. VB assisted with data analysis under the supervision of DI and contributed to planarian maintenance. JHH assisted with BMC analysis in R. ZM co-maintained the screening platform and assisted with screening. CR assisted with screening, performed Ellman assays, and maintained planarian laboratory stocks. EMC contributed to experimental design and coordination of the experiments, performed screening experiments and contributed to data analysis and interpretation. DI and EMC wrote the first draft of the manuscript. All authors contributed to the final version of the manuscript.

## Acknowledgments

The authors thank Dr. Pamela Lein and Dr. Theodor Slotkin for discussions and Dr. Cynthia Rider for comments on the manuscript.

## 8 Data Availability Statement

The datasets generated for this study can be found in the Dryad Digital Repository: https://doi.org/10.5061/dryad.hdr7sqvkr.

## References

1. Adler, M., Manley, H. A., Purcell, A. L., Deshpande, S. S., Hamilton, T. A., Kan, R. K., et al. (2004). Reduced acetylcholine receptor density, morphological remodeling, and butyrylcholinesterase activity can sustain muscle function in acetylcholinesterase knockout mice. Muscle Nerve 30, 317–27. doi:10.1002/mus.20099.

2. Akiyama, Y., Agata, K., and Inoue, T. (2015). Spontaneous behaviors and wall-curvature lead to apparent wall preference in planarian. PLoS One 10, e0142214. Available at: https://journals.plos.org/plosone/article?id=10.1371/journal.pone.0142214 [Accessed January 11, 2019].

3. Aldridge, J. E., Levin, E. D., Seidler, F. J., and Slotkin, T. A. (2005). Developmental exposure of rats to chlorpyrifos leads to behavioral alterations in adulthood, involving serotonergic mechanisms and resembling animal models of depression. Environ. Health Perspect. 113, 527–31. doi:10.1289/EHP.7867.

4. Ando, M., and Wakamatsu, K. (1982). Inhibitory effect of acephate (N-acetyl O, S-dimethyl thiophosphoramide) on serum cholinesterase--effect of acephate on cholinesterase. J. Toxicol. Sci. 7, 185–192. doi:10.2131/JTS.7.185.

5. Atwood, D., and Paisley-Jones, C. (2017). Pesticides Industry Sales and Usage 2008 - 2012 Market Estimates. Washington, DC Available at: https://www.epa.gov/sites/production/files/2017-01/documents/pesticides-industry-sales-usage-2016_0.pdf [Accessed September 10, 2017].

6. Birkholz, T. R., and Beane, W. S. (2017). The planarian TRPA1 homolog mediates extraocular behavioral responses to near-ultraviolet light. J. Exp. Biol. 220, 2616–2625. doi:10.1242/jeb.152298.

7. Brown, D. D. R., and Pearson, B. J. (2015). “One FISH, dFISH, three FISH: sensitive methods of whole-mount fluorescent in situ hybridization in freshwater planarians,” in In Situ Hybridization Methods, ed. G. Hauptmann (New York: Springer Science), 127–150. doi:10.1007/978-1-4939-2303-8.

8. Brown, H. M., Ito, H., and Ogden, T. E. (1968). Spectral sensitivity of the planarian ocellus. J. Gen. Physiol. 51, 255. doi:10.1085/JGP.51.2.255.

9. Burke, R. D., Todd, S. W., Lumsden, E., Mullins, R. J., Mamczarz, J., Fawcett, W. P., et al. (2017). Developmental neurotoxicity of the organophosphorus insecticide chlorpyrifos: from clinical findings to preclinical models and potential mechanisms. J. Neurochem. 142, 162–177. doi:10.1111/jnc.14077.

10. Bushnell, P. J., Pope, C. N., and Padilla, S. (1993). Behavioral and neurochemical effects of acute chlorpyrifos in rats: tolerance to prolonged inhibition of cholinesterase. J. Pharmacol. Exp. Ther. 266.

11. Buttarelli, F. R., Pellicano, C., and Pontieri, F. E. (2008). Neuropharmacology and behavior in planarians: Translations to mammals. Comp. Biochem. Physiol. Part C Toxicol. Pharmacol. 147, 399–408. doi:10.1016/j.cbpc.2008.01.009.

12. Buttarelli, F. R., Pontieri, F. E., Margotta, V., and Palladini, G. (2000). Acetylcholine/dopamine interaction in planaria. Comp. Biochem. Physiol. C. Toxicol. Pharmacol. 125, 225–31. doi:10.1016/S0742-8413(99)00111-5.

13. Buttarelli, F. R., Pontieri, F. E., Margotta, V., and Palladini, G. (2002). Cannabinoid-induced stimulation of motor activity in planaria through an opioid receptor-mediated mechanism. Prog. Neuro-Psychopharmacology Biol. Psychiatry 26, 65–68. doi:10.1016/S0278-5846(01)00230-5.

14. Carr, R. L., Graves, C. A., Magnum, L. C., Nail, C. A., and Ross, M. K. (2014). Low level chlorpyrifos exposure increases anandamide accumulation in juvenile rat brain in the absence of brain cholinesterase inhibition. Neurotoxicology 43, 82–89. doi:10.1016/J.NEURO.2013.12.009.

15. Casida, J. E., and Quistad, G. B. (2004). Organophosphate toxicology: safety aspects of nonacetylcholinesterase secondary targets. Chem. Res. Toxicol. 17, 983–998. doi:10.1021/TX0499259.

16. Cassel, J. C., and Jeltsch, H. (1995). Serotonergic modulation of cholinergic function in the central nervous system: cognitive implications. Neuroscience 69, 1–41. doi:10.1016/0306-4522(95)00241-A.

17. Caughlan, A., Newhouse, K., Namgung, U., and Xia, Z. (2004). Chlorpyrifos induces apoptosis in rat cortical neurons that is regulated by a balance between p38 and ERK/JNK MAP kinases. Toxicol. Sci. 78, 125–34. doi:10.1093/toxsci/kfh038.

18. Cebrià, F. (2007). Regenerating the central nervous system: how easy for planarians! *Dev*. Genes Evol. 217, 733–48. doi:10.1007/s00427-007-0188-6.

19. Chatonnet, F., Boudinot, É., Chatonnet, A., Taysse, L., Daulon, S., Champagnat, J., et al. (2003). Respiratory survival mechanisms in acetylcholinesterase knockout mouse. Eur. J. Neurosci. 18, 1419–1427. doi:10.1046/J.1460-9568.2003.02867.X.

20. Chavarría, A. P., Swartzwelder, J. C., Villarejos, V. M., Kotcher, E., and Arguedas, J. (1969). Dichlorvos, an effective broad-spectrum anthelmintic. Am. J. Trop. Med. Hyg. 18, 907–911. doi:10.4269/ajtmh.1969.18.907.

21. Cochet-Escartin, O., Carter, J. A., Chakraverti-Wuerthwein, M., Sinha, J., and Collins, E. M. S. (2016). Slo1 regulates ethanol-induced scrunching in freshwater planarians. Phys. Biol. 13, 1–12. doi:10.1088/1478-3975/13/5/055001.

22. Cochet-Escartin, O., Mickolajczk, K. J., and Collins, E.-M. S. (2015). Scrunching: a novel escape gait in planarians. Phys. Biol. 12, 055001. doi:10.1088/1478-3975/12/5/056010.

23. Cole, R. D., Anderson, G. L., and Williams, P. L. (2004). The nematode Caenorhabditis elegans as a model of organophosphate-induced mammalian neurotoxicity. Toxicol. Appl. Pharmacol. 194, 248–56. doi:10.1016/j.taap.2003.09.013.

24. Costa, L. G. (2018). Organophosphorus compounds at 80: Some old and new issues. Toxicol. Sci. 162, 24–35. doi:10.1093/toxsci/kfx266.

25. Crumpton, T. L., Seidler, F. J., and Slotkin, T. A. (2000). Is oxidative stress involved in the developmental neurotoxicity of chlorpyrifos? Dev. Brain Res. 121, 189–195. doi:10.1016/S0165-3806(00)00045-6.

26. Dam, K., Seidler, F. J., and Slotkin, T. A. (2000). Chlorpyrifos exposure during a critical neonatal period elicits gender-selective deficits in the development of coordination skills and locomotor activity. Brain Res. Dev. Brain Res. 121, 179–87. Available at: http://www.ncbi.nlm.nih.gov/pubmed/10876030 [Accessed June 8, 2018].

27. dos Santos, A. A., Naime, A. A., de Oliveira, J., Colle, D., dos Santos, D. B., Hort, M. A., et al. (2016). Long-term and low-dose malathion exposure causes cognitive impairment in adult mice: evidence of hippocampal mitochondrial dysfunction, astrogliosis and apoptotic events. Arch. Toxicol. 90, 647–660. doi:10.1007/S00204-015-1466-0.

28. Dunkel, J., Talbot, J., and Schötz, E.-M. (2011). Memory and obesity affect the population dynamics of asexual freshwater planarians. Phys. Biol. 8, 026003. doi:10.1088/1478-3975/8/2/026003.

29. Duysen, E. G., Stribley, J. A., Fry, D. L., Hinrichs, S. H., and Lockridge, O. (2002). Rescue of the acetylcholinesterase knockout mouse by feeding a liquid diet; phenotype of the adult acetylcholinesterase deficient mouse. Brain Res. Dev. Brain Res. 137, 43–54. doi:10.1016/S0165-3806(02)00367-X.

30. Dvergsten, C., and Meeker, R. B. (1994). Muscarinic cholinergic receptor regulation and acetylcholinesterase inhibition in response to insecticide exposure during development. Int. J. Dev. Neurosci. 12, 63–75. doi:10.1016/0736-5748(94)90097-3.

31. Ellman, G. L., Courtney, K. D., Andres, V., and Featherstone, R. M. (1961). A new and rapid colorimetric determination of acetylcholinesterase activity. Biochem. Pharmacol. 7, 88–95. doi:10.1016/0006-2952(61)90145-9.

32. EUROSTAT (2016). *Agriculture, forestry and fishery statistics - 2016 edition*., eds. R. Forti and M. Henrard Luxembourg, Belgium: European Union doi:10.2785/917017.

33. Feldhaus, J. M., Feldhaus, A. J., Ace, L. N., and Pope, C. N. (1998). “Interactive effects of pesticide mixtures on the neurobehavioral responses and AChE levels of planaria,” in Environmental Toxicology and Risk Assessment: Seventh Volume, ASTM STP 1333, eds. E. E. Little, A. J. DeLonay, and B. M. Greenberg (American Society for Testing and Materials), 140–150. Available at: https://books.google.com/books?id=6EPcVPX7AZ8C&pg=PA147&lpg=PA147&dq=Carbaryl+planaria&source=bl&ots=5qUNGaWATY&sig=x6DRBpnBCR0aTuTNlWeGTgPEqMo&hl=en&sa=X&ved=0ahUKEwiK6OjJ4-LZAhUM9GMKHao_AJcQ6AEIPzAD#v=onepage&q=Carbaryl_planaria&f=false [Accessed March 10, 2018].

34. Flaskos, J. (2012). The developmental neurotoxicity of organophosphorus insecticides: a direct role for the oxon metabolites. Toxicol. Lett. 209, 86–93. doi:10.1016/j.toxlet.2011.11.026.

35. Flaskos, J. (2014). The neuronal cytoskeleton as a potential target in the developmental neurotoxicity of organophosphorothionate insecticides. Basic Clin. Pharmacol. Toxicol. 115, 201–8. doi:10.1111/bcpt.12204.

36. Fortunato, J. J., Feier, G., Vitali, A. M., Petronilho, F. C., Dal-Pizzol, F., and Quevedo, J. (2006). Malathion-induced oxidative stress in rat brain regions. Neurochem. Res. 31, 671–678. doi:10.1007/S11064-006-9065-3/FIGURES/4.

37. Gearhart, D. A., Sickles, D. W., Buccafusco, J. J., Prendergast, M. A., and Terry, A. V. (2007). Chlorpyrifos, chlorpyrifos-oxon, and diisopropylfluorophosphate inhibit kinesin-dependent microtubule motility. Toxicol. Appl. Pharmacol. 218, 20–29. doi:10.1016/J.TAAP.2006.10.008.

38. González-Alzaga, B., Lacasaña, M., Aguilar-Garduño, C., Rodríguez-Barranco, M., Ballester, F., Rebagliato, M., et al. (2014). A systematic review of neurodevelopmental effects of prenatal and postnatal organophosphate pesticide exposure. Toxicol. Lett. 230, 104–121. doi:10.1016/j.toxlet.2013.11.019.

39. Grigoryan, H., Schopfer, L. M., Thompson, C. M., Terry, A. V., Masson, P., and Lockridge, O. (2008). Mass spectrometry identifies covalent binding of soman, sarin, chlorpyrifos oxon, diisopropyl fluorophosphate, and FP-biotin to tyrosines on tubulin: a potential mechanism of long term toxicity by organophosphorus agents. Chem. Biol. Interact. 175, 180. doi:10.1016/J.CBI.2008.04.013.

40. Gu, Z., Gu, L., Eils, R., Schlesner, M., and Brors, B. (2014). circlize Implements and enhances circular visualization in R. Bioinformatics 30, 2811–2812. doi:10.1093/BIOINFORMATICS/BTU393.

41. Guizzetti, M., Pathak, S., Giordano, G., and Costa, L. G. (2005). Effect of organophosphorus insecticides and their metabolites on astroglial cell proliferation. Toxicology 215, 182–90. doi:10.1016/j.tox.2005.07.004.

42. Hagstrom, D., Cochet-Escartin, O., and Collins, E.-M. S. (2016). Planarian brain regeneration as a model system for developmental neurotoxicology. Regeneration 3, 65–77. doi:10.1002/reg2.52.

43. Hagstrom, D., Cochet-Escartin, O., Zhang, S., Khuu, C., and Collins, E.-M. S. (2015). Freshwater planarians as an alternative animal model for neurotoxicology. Toxicol. Sci. 147, 270–285. doi:10.1093/toxsci/kfv129.

44. Hagstrom, D., Hirokawa, H., Zhang, L., Radić, Z., Taylor, P., and Collins, E.-M. S. (2017a). Planarian cholinesterase: in vitro characterization of an evolutionarily ancient enzyme to study organophosphorus pesticide toxicity and reactivation. Arch. Toxicol. 91, 2837–2847. doi:10.1007/s00204-016-1908-3.

45. Hagstrom, D., Hirokawa, H., Zhang, L., Radić, Z., Taylor, P., and Collins, E.-M. S. (2017b). Planarian cholinesterase: in vitro characterization of an evolutionarily ancient enzyme to study organophosphorus pesticide toxicity and reactivation. Arch. Toxicol. 91, 2837–2847. doi:10.1007/s00204-016-1908-3.

46. Hagstrom, D., Truong, L., Zhang, S., Tanguay, R., and Collins, E.-M. S. (2019). Comparative analysis of zebrafish and planarian model systems for developmental neurotoxicity screens using an 87-compound library. Toxicol. Sci. 167, 15–25. doi:10.1093/toxsci/kfy180.

47. Hagstrom, D., Zhang, S., Ho, A., Tsai, E. S., Radić, Z., Jahromi, A., et al. (2018). Planarian cholinesterase: molecular and functional characterization of an evolutionarily ancient enzyme to study organophosphorus pesticide toxicity. Arch. Toxicol. 92, 1161–1176. doi:10.1007/s00204-017-2130-7.

48. Howard, M. D., and Pope, C. N. (2002). In vitro effects of chlorpyrifos, parathion, methyl parathion and their oxons on cardiac muscarinic receptor binding in neonatal and adult rats. Toxicology 170, 1–10. doi:10.1016/S0300-483X(01)00498-X.

49. Hsieh, J. H., Ryan, K., Sedykh, A., Lin, J. A., Shapiro, A. J., Parham, F., et al. (2019). Application of benchmark concentration (BMC) analysis on zebrafish data: A new perspective for quantifying toxicity in alternative animal models. Toxicol. Sci. 167, 282–292. doi:10.1093/toxsci/kfy258.

50. Hyman, L. H. (1919). Physiological studies on Planaria III. Oxygen consumption in relation to age (size) differences. Biol. Bull. 37, 388–403. doi:10.2307/1536374.

51. Inoue, T., Hoshino, H., Yamashita, T., Shimoyama, S., and Agata, K. (2015). Planarian shows decision-making behavior in response to multiple stimuli by integrative brain function. Zool. Lett., 1–7. Available at: http://zoologicalletters.biomedcentral.com/articles/10.1186/s40851-014-0010-z [Accessed November 4, 2015].

52. Inoue, T., Yamashita, T., and Agata, K. (2014). Thermosensory signaling by TRPM is processed by brain serotonergic neurons to produce planarian thermotaxis. J. Neurosci. 34, 15701–14. doi:10.1523/JNEUROSCI.5379-13.2014.

53. Ireland, D., Bochenek, V., Chaiken, D., Rabeler, C., Onoe, S., Soni, A., et al. (2020). Dugesia japonica is the best suited of three planarian species for high-throughput toxicology screening. Chemosphere 253, 126718. doi:10.1016/j.chemosphere.2020.126718.

54. Jansen, K. L., Cole, T. B., Park, S. S., Furlong, C. E., and Costa, L. G. (2009). Paraoxonase 1 (PON1) modulates the toxicity of mixed organophosphorus compounds. Toxicol. Appl. Pharmacol. 236, 142–53. doi:10.1016/j.taap.2009.02.001.

55. Jenkins, M. M. (1959). Respiration rates in Planarians I. The use of the Warburg respirometer in determining oxygen consumption. Proc. Okla. Acad. Sci., 35–40.

56. Jett, D. A., Hill, E. F., Fernando, J. C., Eldefrawi, M. E., and Eldefrawi, A. T. (1993). Down-regulation of muscarinic receptors and the m3 subtype in white-footed mice by dietary exposure to parathion. J. Toxicol. Environ. Health 39, 395–415. doi:10.1080/15287399309531760.

57. Jiang, W., Duysen, E. G., Hansen, H., Shlyakhtenko, L., Schopfer, L. M., and Lockridge, O. (2010). Mice treated with chlorpyrifos or chlorpyrifos oxon have organophosphorylated tubulin in the brain and disrupted microtubule structures, suggesting a role for tubulin in neurotoxicity associated with exposure to organophosphorus agents. Toxicol. Sci. 115, 183–193. doi:10.1093/toxsci/kfq032.

58. Kaur, J., and Singh, P. K. (2020). Enzyme-based optical biosensors for organophosphate class of pesticide detection. Phys. Chem. Chem. Phys. 22, 15105–15119. doi:10.1039/D0CP01647K.

59. Koenig, J. A., Chen, C. A., and Shih, T. M. (2020). Development of a larval zebrafish model for acute organophosphorus nerve agent and pesticide exposure and therapeutic evaluation. Toxics 8, 1–11. doi:10.3390/toxics8040106.

60. Lein, P. J., and Fryer, A. D. (2005). Organophosphorus insecticides induce airway hyperreactivity by decreasing neuronal M2 muscarinic receptor function independent of acetylcholinesterase inhibition. Toxicol. Sci. 83, 166–176. doi:10.1093/TOXSCI/KFI001.

61. Levin, E. D., Swain, H. A., Donerly, S., and Linney, E. (2004). Developmental chlorpyrifos effects on hatchling zebrafish swimming behavior. Neurotoxicol. Teratol. 26, 719–723. doi:10.1016/j.ntt.2004.06.013.

62. Levy, R., and Miller, T. W. (1978). Tolerance of the planarian Dugesia dorotocephala to high concentrations of pesticides and growth regulators. Entomophaga 23, 31–34. doi:10.1007/BF02371988.

63. Levy, S. J., and Perron, M. M. (2016). Amendment to: Profenofos: Human Health Draft Risk Assessment (DRA) for Registration Review.

64. Lionetto, M. G., Caricato, R., Calisi, A., Giordano, M. E., and Schettino, T. (2013). Acetylcholinesterase as a biomarker in environmental and occupational medicine: New insights and future perspectives. Biomed Res. Int. 2013, 1–8. doi:10.1155/2013/321213.

65. Liu, J., Parsons, L., and Pope, C. (2013). Comparative effects of parathion and chlorpyrifos on extracellular endocannabinoid levels in rat hippocampus: Influence on cholinergic toxicity. Toxicol. Appl. Pharmacol. 272, 608–615. doi:10.1016/j.taap.2013.07.025.

66. Liu, X. Y., Zhang, Q. P., Li, S. B., Mi, P., Chen, D. Y., Zhao, X., et al. (2018). Developmental toxicity and neurotoxicity of synthetic organic insecticides in zebrafish (Danio rerio): A comparative study of deltamethrin, acephate, and thiamethoxam. Chemosphere 199, 16–25. doi:10.1016/J.CHEMOSPHERE.2018.01.176.

67. Malinowski, P. T., Cochet-Escartin, O., Kaj, K. J., Ronan, E., Groisman, A., Diamond, P. H., et al. (2017). Mechanics dictate where and how freshwater planarians fission. Proc. Natl. Acad. Sci. U. S. A. 114, 10888–10893. doi:10.1073/pnas.1700762114.

68. Mamczarz, J., Pescrille, J. D., Gavrushenko, L., Burke, R. D., Fawcett, W. P., DeTolla, L. J., et al. (2016). Spatial learning impairment in prepubertal guinea pigs prenatally exposed to the organophosphorus pesticide chlorpyrifos: Toxicological implications. Neurotoxicology 56, 17–28. doi:10.1016/j.neuro.2016.06.008.

69. Moser, V. C. (1995). Comparisons of the acute effects of cholinesterase inhibitors using a neurobehavioral screening battery in rats. Neurotoxicol. Teratol. 17, 617–625. doi:10.1016/0892-0362(95)02002-0.

70. Muñoz-Quezada, M. T., Lucero, B. A., Barr, D. B., Steenland, K., Levy, K., Ryan, P. B., et al. (2013). Neurodevelopmental effects in children associated with exposure to organophosphate pesticides: a systematic review. Neurotoxicology 39, 158–168. doi:10.1016/j.neuro.2013.09.003.

71. Nishimura, K., Kitamura, Y., Taniguchi, T., and Agata, K. (2010). Analysis of motor function modulated by cholinergic neurons in planarian Dugesia japonica. Neuroscience 168, 18–30. doi:10.1016/j.neuroscience.2010.03.038.

72. Nishimura, O., Hosoda, K., Kawaguchi, E., Yazawa, S., Hayashi, T., Inoue, T., et al. (2015). Unusually large number of mutations in asexually reproducing clonal planarian Dugesia japonica. PLoS One 10, 1–23. doi:10.1371/journal.pone.0143525.

73. Osuma, E. A., Riggs, D. W., Gibb, A. A., and Hill, B. G. (2018). High throughput measurement of metabolism in planarians reveals activation of glycolysis during regeneration. Regeneration 5, 78. doi:10.1002/REG2.95.

74. Pagán, O. R. (2014). The first brain: the neuroscience of planarians. Oxford University Press.

75. Pagán, O. R., Montgomery, E., Deats, S., Bach, D., and Baker, D. (2015). Evidence of nicotine-induced, curare-insensitive, behavior in planarians. Neurochem. Res. 40, 2087–2090. doi:10.1007/s11064-015-1512-6.

76. Paskin, T. R., Jellies, J., Bacher, J., and Beane, W. S. (2014). Planarian phototactic assay reveals differential behavioral responses based on wavelength. PLoS One 9, e114708. doi:10.1371/journal.pone.0114708.

77. Peeples, E. S., Schopfer, L. M., Duysen, E. G., Spaulding, R., Voelker, T., Thompson, C. M., et al. (2005). Albumin, a new biomarker of organophosphorus toxicant exposure, identified by mass spectrometry. Toxicol. Sci. 83, 303–312. doi:10.1093/TOXSCI/KFI023.

78. Peter, J. V., Sudarsan, T. I., and Moran, J. L. (2014). Clinical features of organophosphate poisoning: A review of different classification systems and approaches. Indian J. Crit. Care Med. 18, 735. doi:10.4103/0972-5229.144017.

79. Poirier, L., Brun, L., Jacquet, P., Lepolard, C., Armstrong, N., Torre, C., et al. (2017). Enzymatic degradation of organophosphorus insecticides decreases toxicity in planarians and enhances survival. Sci. Rep. 7, 15194. doi:10.1038/s41598-017-15209-8.

80. Pope, C., Karanth, S., and Liu, J. (2005a). Pharmacology and toxicology of cholinesterase inhibitors: uses and misuses of a common mechanism of action. Environ. Toxicol. Pharmacol. 19, 433–46. doi:10.1016/j.etap.2004.12.048.

81. Pope, C., Karanth, S., and Liu, J. (2005b). Pharmacology and toxicology of cholinesterase inhibitors: uses and misuses of a common mechanism of action. Environ. Toxicol. Pharmacol. 19, 433–446. doi:10.1016/j.etap.2004.12.048.

82. Pope, C. N. (1999). Organophosphorus pesticides: do they all have the same mechanism of toxicity? J. Toxicol. Environ. Heal. Part B Crit. Rev. 2, 161–181. doi:10.1080/109374099281205.

83. Prendergast, M. A., Self, R. L., Smith, K. J., Ghayoumi, L., Mullins, M. M., Butler, T. R., et al. (2007). Microtubule-associated targets in chlorpyrifos oxon hippocampal neurotoxicity. Neuroscience 146, 330. doi:10.1016/J.NEUROSCIENCE.2007.01.023.

84. Proskocil, B. J., Bruun, D. A., Thompson, C. M., Fryer, A. D., and Lein, P. J. (2010). Organophosphorus pesticides decrease M2 muscarinic receptor function in guinea pig airway nerves via indirect mechanisms. PLoS One 5, e10562. doi:10.1371/JOURNAL.PONE.0010562.

85. R Core Team (2016). R: A language environment for statistical computing. Vienna, Austria Available at: https://www.r-project.org.

86. Rajini, P. S., Melstrom, P., and Williams, P. L. (2008). A comparative study on the relationship between various toxicological endpoints in Caenorhabditis elegans exposed to organophosphorus insecticides. J. Toxicol. Environ. Heal. Part A Curr. Issues 71, 1043–1050. doi:10.1080/15287390801989002.

87. Rauh, V. A., Perera, F. P., Horton, M. K., Whyatt, R. M., Bansal, R., Hao, X., et al. (2012). Brain anomalies in children exposed prenatally to a common organophosphate pesticide. Proc. Natl. Acad. Sci. U. S. A. 109, 7871–6. doi:10.1073/pnas.1203396109.

88. Rauh, V., Arunajadai, S., Horton, M., Perera, F., Hoepner, L., Barr, D. B., et al. (2011). Seven-year neurodevelopmental scores and prenatal exposure to chlorpyrifos, a common agricultural pesticide. Environ. Health Perspect. 119, 1196–1201. doi:10.1289/ehp.1003160.

89. Rawls, S. M., Patil, T., Tallarida, C. S., Baron, S., Kim, M., Song, K., et al. (2011). Nicotine behavioral pharmacology: Clues from planarians. Drug Alcohol Depend. 118, 274–279. doi:10.1016/j.drugalcdep.2011.04.001.

90. Razwiedani, L. L., and Rautenbach, P. (2017). Epidemiology of Organophosphate Poisoning in the Tshwane District of South Africa. Environ. Health Insights 11, 1178630217694149. doi:10.1177/1178630217694149.

91. Ribeiro, P., El-Shehabi, F., and Patocka, N. (2005). Classical transmitters and their receptors in flatworms. Parasitology 131, S19–40. doi:10.1017/S0031182005008565.

92. Richendrfer, H., and Creton, R. (2015). Chlorpyrifos and malathion have opposite effects on behaviors and brain size that are not correlated to changes in AChE activity. Neurotoxicology 49, 50–58. doi:10.1016/j.neuro.2015.05.002.

93. Ritz, C., Baty, F., Streibig, J. C., and Gerhard, D. (2015). Dose-response analysis Using R. PLoS One 10, e0146021. doi:10.1371/JOURNAL.PONE.0146021.

94. Rompolas, P., Azimzadeh, J., Marshall, W. F., and King, S. M. (2013). Analysis of ciliary assembly and function in planaria. Methods Enzymol. 525, 245–264. doi:10.1016/B978-0-12-397944-5.00012-2.

95. Ross, K. G., Currie, K. W., Pearson, B. J., and Zayas, R. M. (2017). Nervous system development and regeneration in freshwater planarians. Wiley Interdiscip. Rev. Dev. Biol. 6, 1–26. doi:10.1002/wdev.266.

96. Russom, C. L., LaLone, C. A., Villeneuve, D. L., and Ankley, G. T. (2014). Development of an adverse outcome pathway for acetylcholinesterase inhibition leading to acute mortality. Environ. Toxicol. Chem. 33, 2157–2169. doi:10.1002/etc.2662.

97. Sabry, Z., Ho, A., Ireland, D., Rabeler, C., Cochet-Escartin, O., and Collins, E. M. S. (2019). Pharmacological or genetic targeting of Transient Receptor Potential (TRP) channels can disrupt the planarian escape response. PLoS One 14, e0226104. doi:10.1371/journal.pone.0226104.

98. Sabry, Z., Rabeler, C., Ireland, D., Bayingana, K., and Collins, E. M. S. (2020). Planarian scrunching as a quantitative behavioral readout for noxious stimuli sensing. J. Vis. Exp. 2020, 1–18. doi:10.3791/61549.

99. Sagiv, S. K., Kogut, K., Harley, K., Bradman, A., Morga, N., and Eskenazi, B. (2021). Gestational exposure to organophosphate pesticides and longitudinally assessed behaviors related to Attention-Deficit/Hyperactivity Disorder and executive function. Am. J. Epidemiol. 190, 2420– 2431. doi:10.1093/AJE/KWAB173.

100. Samimi, A., and Last, J. A. (2001). Inhibition of lysyl hydroxylase by malathion and malaoxon. Toxicol. Appl. Pharmacol. 172, 203–209. doi:10.1006/taap.2001.9147.

101. Schmitt, C., McManus, M., Kumar, N., Awoyemi, O., and Crago, J. (2019). Comparative analyses of the neurobehavioral, molecular, and enzymatic effects of organophosphates on embryo-larval zebrafish (Danio rerio). Neurotoxicol. Teratol. 73, 67–75. doi:10.1016/J.NTT.2019.04.002.

102. Shelton, J. F., Geraghty, E. M., Tancredi, D. J., Delwiche, L. D., Schmidt, R. J., Ritz, B., et al. (2014). Neurodevelopmental disorders and prenatal residential proximity to agricultural pesticides: the CHARGE study. Environ. Health Perspect. 122, 1103–9. doi:10.1289/ehp.1307044.

103. Shettigar, N., Joshi, A., Dalmeida, R., Gopalkrishna, R., Chakravarthy, A., Patnaik, S., et al. (2017). Hierarchies in light sensing and dynamic interactions between ocular and extraocular sensory networks in a flatworm. Sci. Adv. 3. doi:10.1126/sciadv.1603025.

104. Slotkin, T. A. (2004). Cholinergic systems in brain development and disruption by neurotoxicants: nicotine, environmental tobacco smoke, organophosphates. Toxicol. Appl. Pharmacol. 198, 132–51. doi:10.1016/j.taap.2003.06.001.

105. Slotkin, T. A. (2006). “Developmental Neurotoxicity of Organophosphates,” in Toxicology of Organophosphate & Carbamate Compounds (Elsevier), 293–314. doi:10.1016/B978-012088523-7/50022-3.

106. Slotkin, T. A., Cousins, M. M., Tate, C. A., and Seidler, F. J. (2001). Persistent cholinergic presynaptic deficits after neonatal chlorpyrifos exposure. Brain Res. 902, 229–243. doi:10.1016/S0006-8993(01)02387-3.

107. Slotkin, T. A., Levin, E. D., and Seidler, F. J. (2006a). Comparative developmental neurotoxicity of organophosphate insecticides: Effects on brain development are separable from systemic toxicity. Environ. Health Perspect. 114, 746–751. doi:10.1289/ehp.8828.

108. Slotkin, T. A., and Seidler, F. J. (2008). Developmental neurotoxicants target neurodifferentiation into the serotonin phenotype: Chlorpyrifos, diazinon, dieldrin and divalent nickel. Toxicol. Appl. Pharmacol. 233, 211–219. doi:https://doi.org/10.1016/j.taap.2008.08.020.

109. Slotkin, T. A., Seidler, F. J., and Fumagalli, F. (2007). Exposure to organophosphates reduces the expression of neurotrophic factors in neonatal rat brain regions: Similarities and differences in the effects of chlorpyrifos and diazinon on the fibroblast growth factor superfamily. Environ. Health Perspect. 115, 909–916. doi:10.1289/EHP.9901.

110. Slotkin, T. A., Skavicus, S., Ko, A., Levin, E. D., and Seidler, F. J. (2019). Perinatal diazinon exposure compromises the development of acetylcholine and serotonin systems. Toxicology 424, 152240. doi:10.1016/j.tox.2019.152240.

111. Slotkin, T. A., Skavicus, S., and Seidler, F. J. (2017). Diazinon and parathion diverge in their effects on development of noradrenergic systems. Brain Res. Bull. 130, 268–273. doi:10.1016/J.BRAINRESBULL.2017.02.004.

112. Slotkin, T. A., Tate, C. A., Ryde, I. T., Levin, E. D., and Seidler, F. J. (2006b). Organophosphate insecticides target the serotonergic system in developing rat brain regions: disparate effects of diazinon and parathion at doses spanning the threshold for cholinesterase inhibition. Environ. Health Perspect. 114, 1542–6. doi:10.1289/EHP.9337.

113. Smulders, C. J. G. M., Bueters, T. J. H., Vailati, S., van Kleef, R. G. D. M., and Vijvergberg, H. P. M. (2004). Block of neuronal nicotinic acetylcholine receptors by organophosphate insecticides. Toxicol. Sci. 82, 545–554. doi:10.1093/TOXSCI/KFH269.

114. Snawder, J. E., and Chambers, J. E. (1993). Osteolathyrogenic effects of malathion in Xenopus embryos. Toxicol. Appl. Pharmacol. 121, 210–216. doi:10.1006/taap.1993.1147.

115. Song, X., Seidler, F. J., Saleh, J. L., Zhang, J., Padilla, S., and Slotkin, T. A. (1997). Cellular mechanisms for developmental toxicity of chlorpyrifos: Targeting the adenylyl cyclase signaling cascade. Toxicol. Appl. Pharmacol. 145, 158–174. doi:10.1006/taap.1997.8171.

116. Taylor, P. (2018). “Anticholinesterase agents,” in Goodman and Gilman’s The Pharmacological Basis of Therapeutics, ed. Laurence L Brunton (San Francisco: McGraw Hill Education), 163– 176.

117. Terry, A. V. J. (2012). Functional consequences of repeated organophosphate exposure: potential non-cholinergic mechanisms. Pharmacol. Ther. 134, 355–65. doi:10.1016/j.pharmthera.2012.03.001.

118. Trevisan, R., Uliano-Silva, M., Pandolfo, P., Franco, J. L., Brocardo, P. S., Santos, A. R. S., et al. (2008). Antioxidant and acetylcholinesterase response to repeated malathion exposure in rat cerebral cortex and hippocampus. Basic Clin. Pharmacol. Toxicol. 102, 365–369. doi:10.1111/j.1742-7843.2007.00182.x.

119. Van Huizen, A. V, Tseng, A.-S., and Beane, W. S. (2017). Methylisothiazolinone toxicity and inhibition of wound healing and regeneration in planaria. Aquat. Toxicol. 191, 226–235. doi: https://doi.org/10.1016/j.aquatox.2017.08.013.

120. Venturini, G., Stocchi, F., Margotta, V., Ruggieri, S., Bravi, D., Bellantuono, P., et al. (1989). A pharmacological study of dopaminergic receptors in planaria. Neuropharmacology 28, 1377– 1382. Available at: http://www.sciencedirect.com/science/article/pii/0028390889900130 [Accessed November 3, 2015].

121. Villar, D., Li, M. H., and Schaeffer, D. J. (1993). Toxicity of organophosphorus pesticides to Dugesia dorotocephala. Bull. Environ. Contam. Toxicol. 51, 80–87. doi:10.1007/BF00201004.

122. Voorhees, J. R., Rohlman, D. S., Lein, P. J., and Pieper, A. A. (2016). Neurotoxicity in Preclinical Models of Occupational Exposure to Organophosphorus Compounds. Front. Neurosci. 10, 590. doi:10.3389/FNINS.2016.00590.

123. Voorhees, J. R., Rohlman, D. S., Lein, P. J., and Pieper, A. A. (2017). Neurotoxicity in preclinical models of occupational exposure to organophosphorus compounds. Front. Neurosci. 10, 590. doi:10.3389/FNINS.2016.00590.

124. Yang, D., Howard, A., Bruun, D., Ajua-Alemanj, M., Pickart, C., and Lein, P. J. (2008). Chlorpyrifos and chlorpyrifos-oxon inhibit axonal growth by interfering with the morphogenic activity of acetylcholinesterase. Toxicol. Appl. Pharmacol. 228, 32–41. doi:10.1016/j.taap.2007.11.005.

125. Yang, D., Lauridsen, H., Buels, K., Chi, L. H., La Du, J., Bruun, D. A., et al. (2011). Chlorpyrifos-oxon disrupts zebrafish axonal growth and motor behavior. Toxicol. Sci. 121, 146–159. doi:10.1093/toxsci/kfr028.

126. Yen, J., Donerly, S., Linney, E. A., Levin, E. D., and Linney, E. A. (2011). Differential acetylcholinesterase inhibition of chlorpyrifos, diazinon and parathion in larval zebrafish. Neurotoxicol. Teratol. 33, 735–741. doi:10.1016/j.ntt.2011.10.004.

127. Zarei, M. H., Soodi, M., Qasemian-Lemraski, M., Jafarzadeh, E., and Taha, M. F. (2015). Study of the chlorpyrifos neurotoxicity using neural differentiation of adipose tissue-derived stem cells. Environ. Toxicol. 31, 1510–1519. doi:10.1002/tox.22155.

128. Zhang, S., Hagstrom, D., Hayes, P., Graham, A., and Collins, E.-M. S. (2019a). Multi-behavioral endpoint testing of an 87-chemical compound library in freshwater planarians. Toxicol. Sci. 167, 26–44. doi:10.1093/toxsci/kfy145.

129. Zhang, S., Ireland, D., Sipes, N. S., Behl, M., and Collins, E.-M. S. (2019b). Screening for neurotoxic potential of 15 flame retardants using freshwater planarians. Neurotoxicol. Teratol. 73, 54–66. doi:10.1016/j.ntt.2019.03.003.

130. Zhang, X., Tong, H., Bi, S., and Pang, Q. (2013). Effects of chlorpyrifos on acute toxicity, mobility and regeneration of planarian Dugesia Japonica. Fresenius Environ. Bull. 22, 2610–2615. Available at: https://www.researchgate.net/publication/286408743_Effects_of_chlorpyrifos_on_acute_toxicity_mobility_and_regeneration_of_planarian_Dugesia_Japonica [Accessed April 26, 2022].

131. Zhu, J., Zhang, X., Xu, Y., Spencer, T. J., Biederman, J., and Bhide, P. G. (2012). Prenatal nicotine exposure mouse model showing hyperactivity, reduced cingulate cortex volume, reduced dopamine turnover, and responsiveness to oral methylphenidate treatment. J. Neurosci. 32, 9410–9418. doi:10.1523/JNEUROSCI.1041-12.2012.

