## Supplementary Material for "Differences in neurotoxic outcomes of organophosphorus pesticides revealed via multi-dimensional screening in adult and regenerating planarians"

### 1 Supplementary Tables

**Supplementary Table 1.** Comparison of potency ranking of OPs between adult and regenerating planarians. The most sensitive BMC (µM) for each OP in either adult or regenerating planarians is listed along with the associated relative rank, with 1 being the most potent.

| OP | $BMC_{adult}$ | Rank | $BMC_{regenerating}$ | Rank |
|---|---|---|---|---|
| Acephate | 229 | 7 | 302 | 7 |
| Chlorpyrifos | 1.7 | 4 | 1.7 | 2 |
| Diazinon | 0.22 | 2 | 9.2 | 4 |
| Dichlorvos | 0.09 | 1 | 0.10 | 1 |
| Malathion | 14 | 6 | 43 | 6 |
| Parathion | 5 | 5 | 17 | 5 |
| Profenofos | 0.29 | 3 | 4.6 | 3 |

### 2 Supplementary Figures

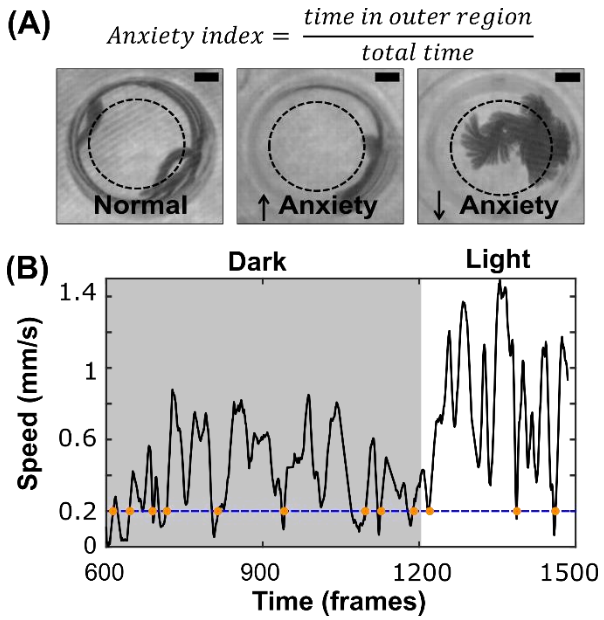


**Supplementary Figure 1. Examples of new endpoints. (A)** Anxiety is measured as the fraction of time a planarian spends in the outer 2/3 of the well (outside the dotted circle). Increased anxiety is associated with less exploration of the well. Scale bar: 2 mm. **(B)** Locomotor bursts (orange dots) are instances where the planarian goes from resting (speed <0.2 mm/s) to not resting (speed >0.2 mm/s). The total cumulative number of locomotor bursts and the ratio of bursts during the blue and 2$^{nd}$ dark phases of the phototaxis assay are quantified.

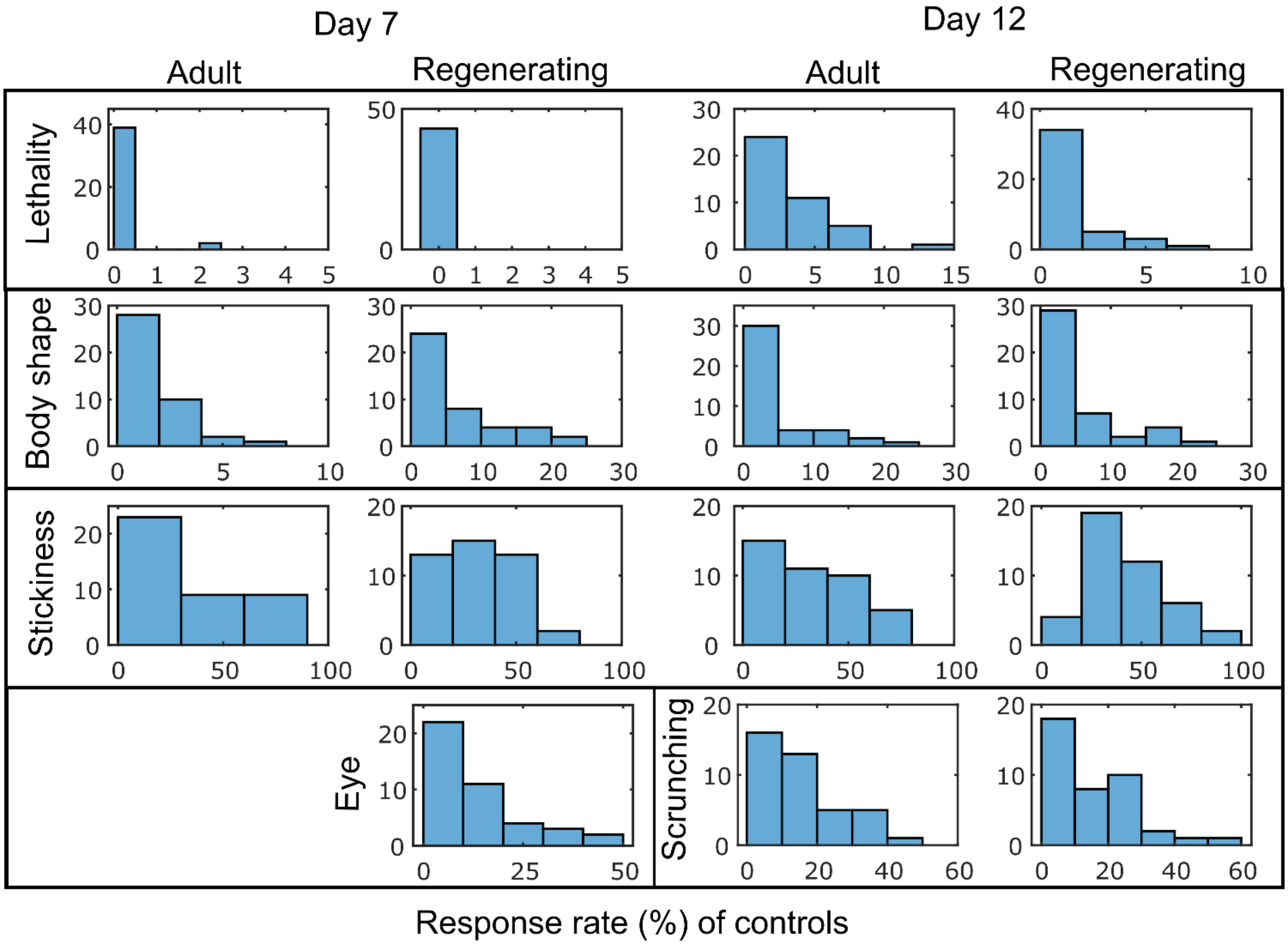


**Supplementary Figure 2. Response rate of vehicle controls in the binary endpoints.** Plots show the distribution of response rates in the vehicle controls for all binary outcome measures.

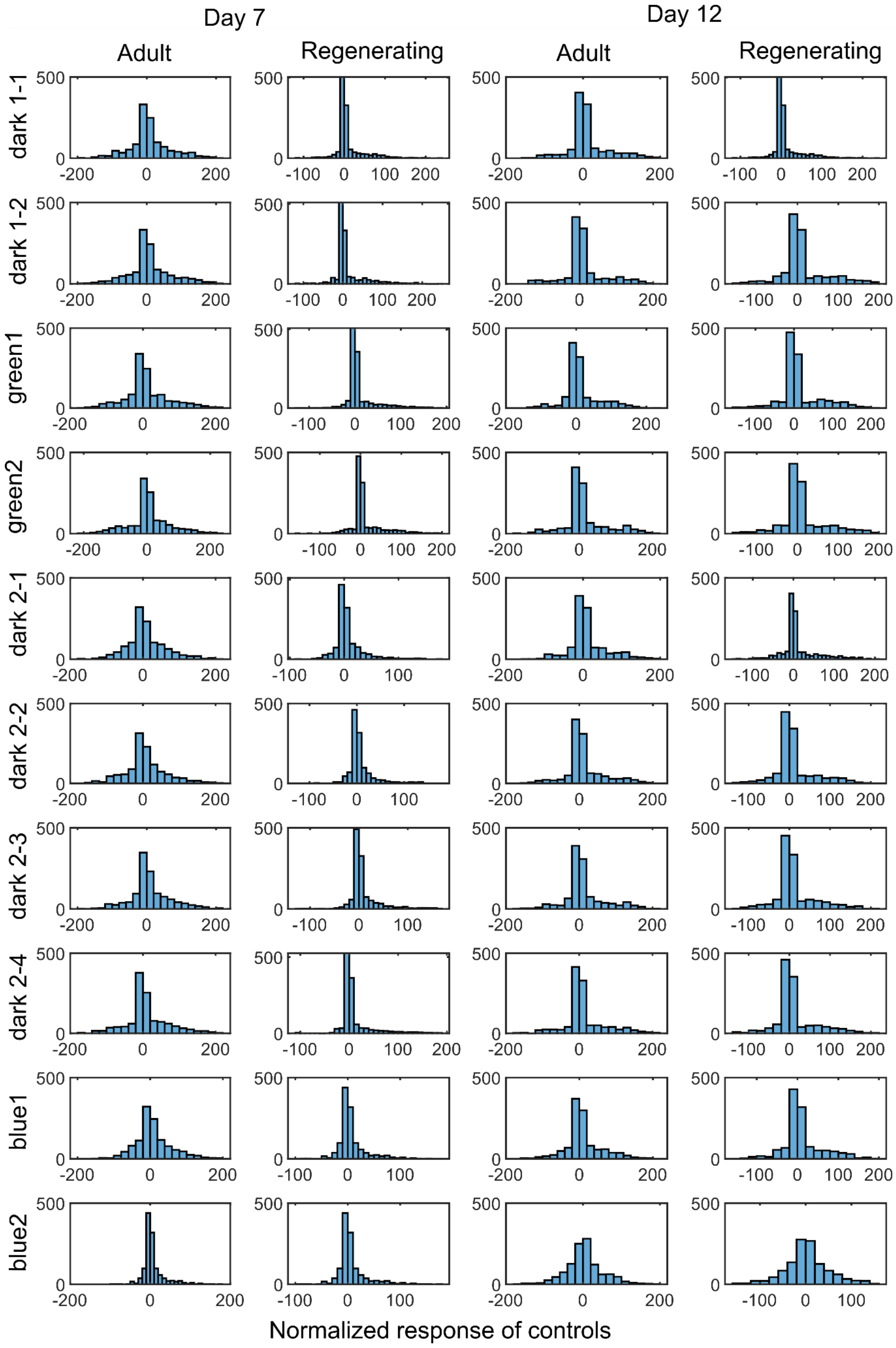
Day 7
Day 12
Adult
Regenerating
Adult
Regenerating
dark 1-1
dark 1-2
green1
green2
dark 2-1
dark 2-2
dark 2-3
dark 2-4
blue1
blue2
Normalized response of controls

**Supplementary Figure 3. Normalized responses of the vehicle controls in the speed endpoints.** Plots show the distribution of normalized responses for each individual vehicle control when normalized by the median of the control population of the respective plate.

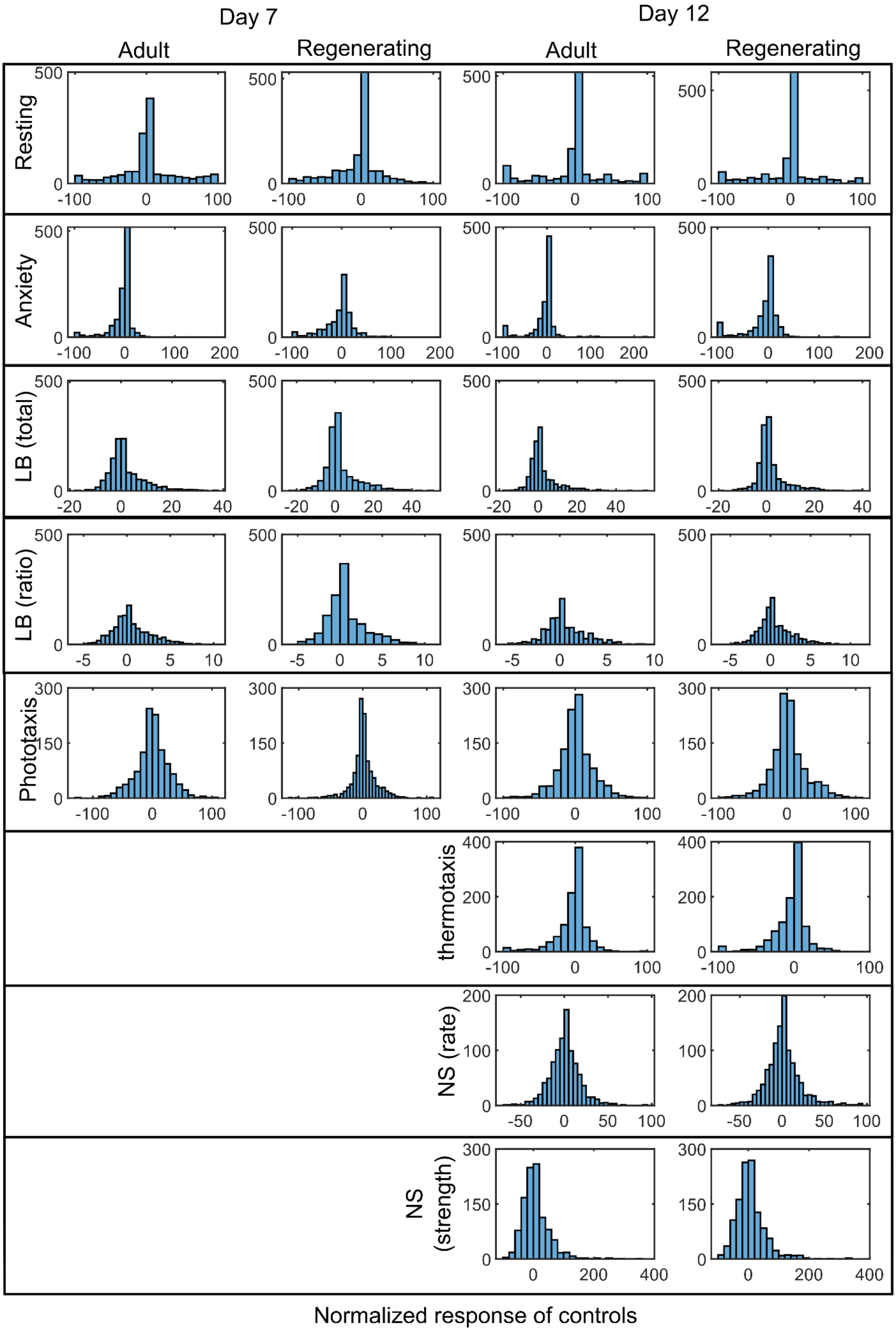
Day 7
Day 12
Adult
Regenerating
Adult
Regenerating
Resting
Anxiety
LB (total)
LB (ratio)
Phototaxis
thermotaxis
NS (rate)
NS (strength)
Normalized response of controls

**Supplementary Figure 4. Normalized responses of the vehicle controls in the remaining continuous endpoints.** Plots show the distribution of normalized responses for each individual vehicle control when normalized by the median of the control population of the respective plate. LB: locomotor bursts; NS: noxious stimuli.

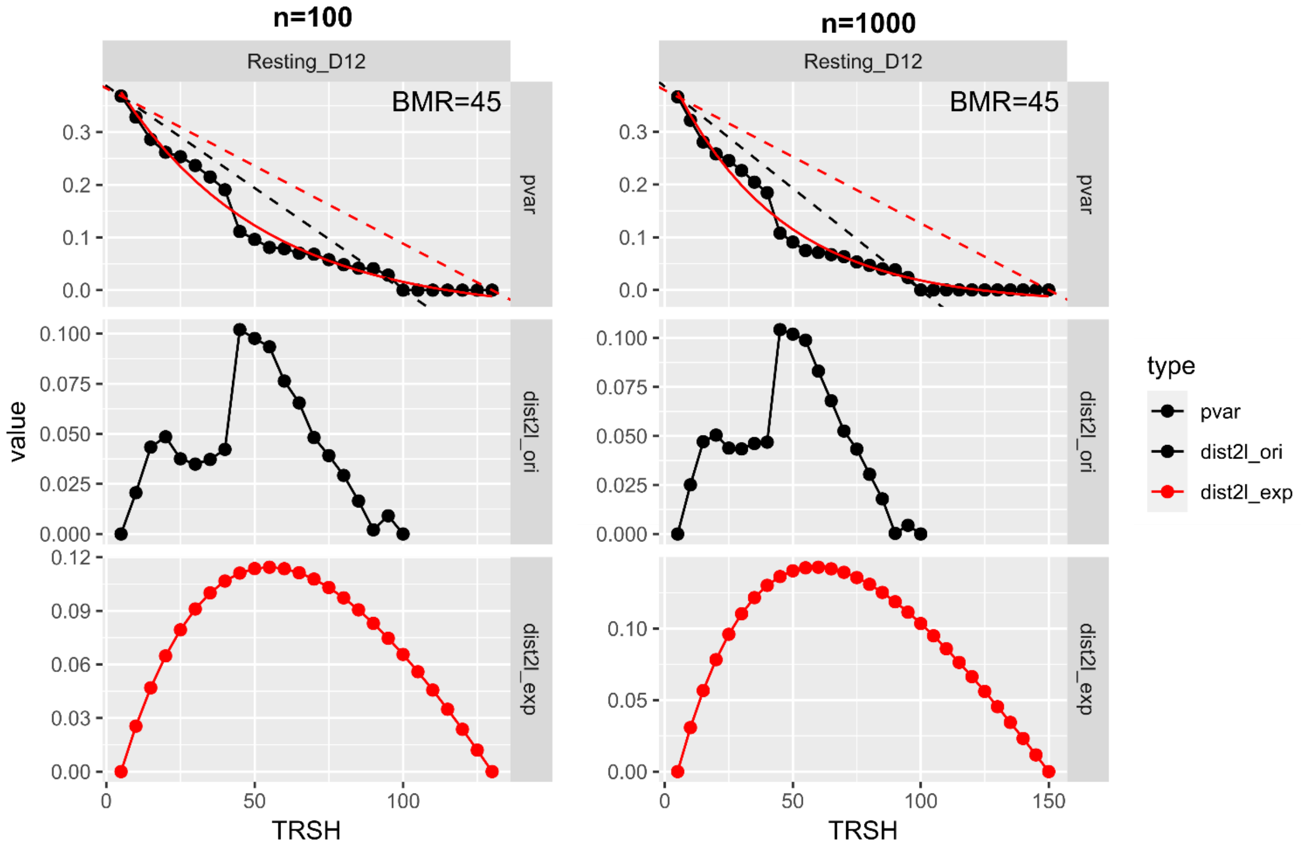


**Supplementary Figure 5. Bootstrapping with n=100 samples gives similar results as n=1000.** Benchmark response (BMR) diagnostic plots were compared when setting the " n_samples" parameter to either 100 or 1000 using the data from the day 12 resting endpoint in adults. The same BMRs were suggested using either number of samples.

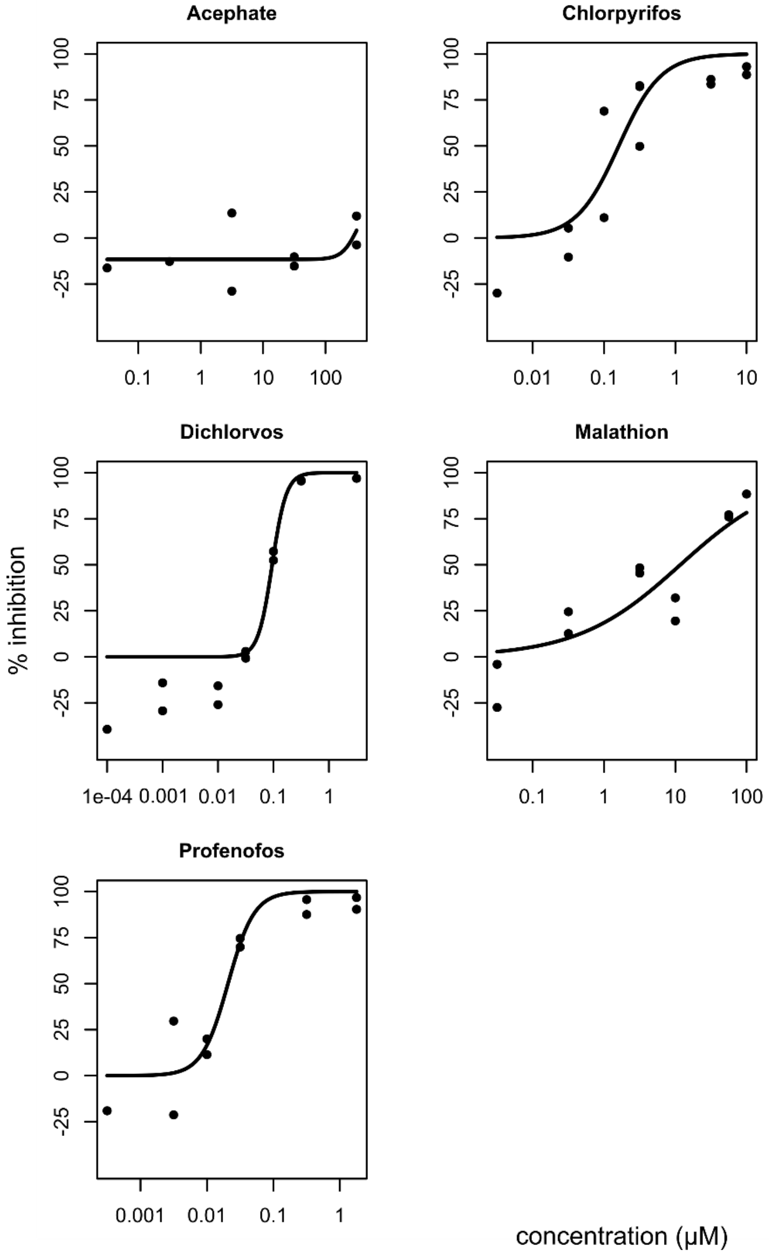


**Supplementary Figure 6. AChE inhibition in adult planarians.** Ellman assays were performed on adult planarians exposed for 12 days to different concentrations of the OPs. Dose response curves were fit with a Hill equation (setting the lower limit to 0 and the upper limit to 100) using the R package drc.

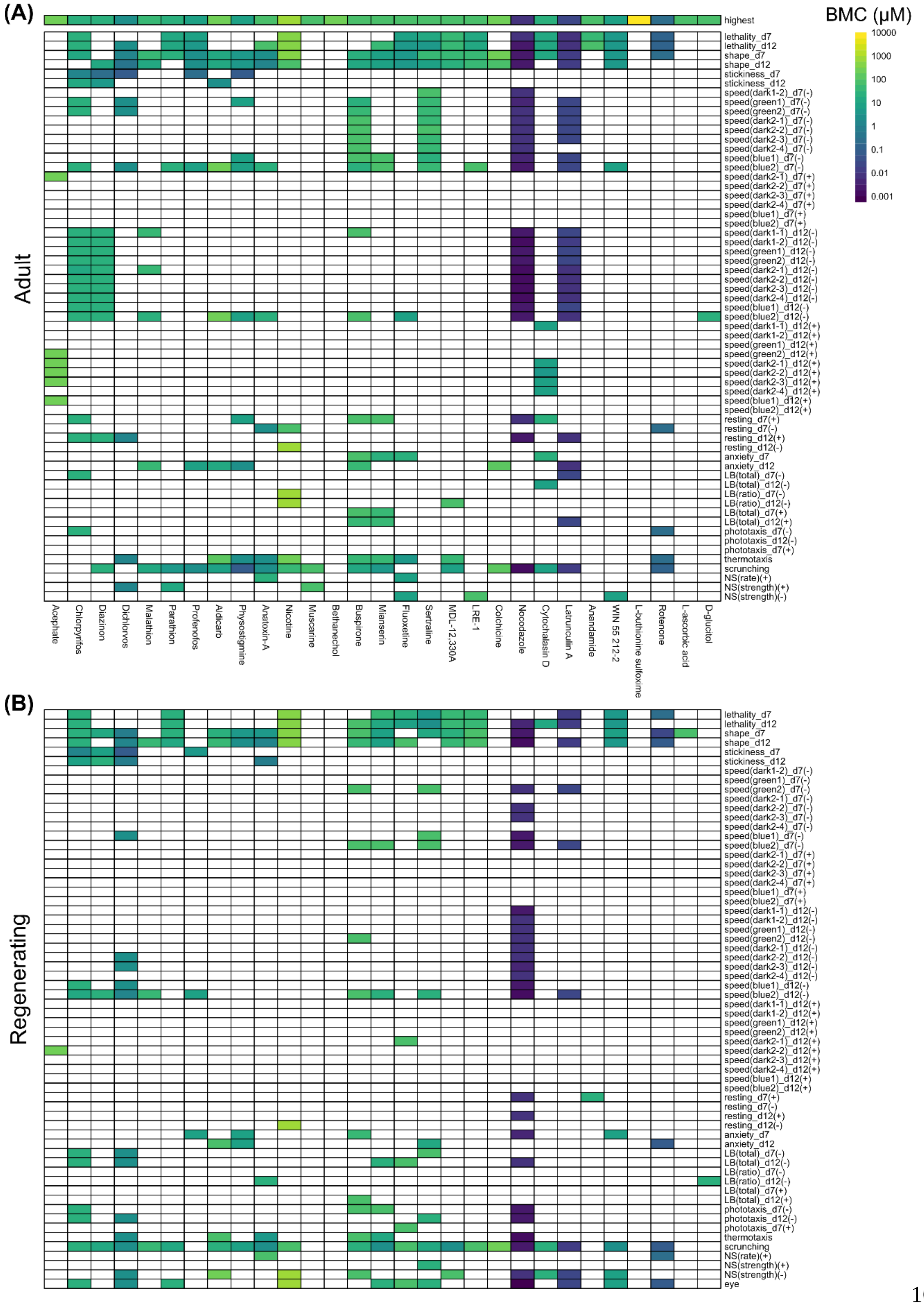
(A)
Adult
BMC (µM)
10000
1000
100
10
1
0.1
0.01
0.001
highest
lethality_d7
lethality_d12
shape_d7
shape_d12
stickiness_d7
stickiness_d12
speed(dark1-2)_d7(-)
speed(green1)_d7(-)
speed(green2)_d7(-)
speed(dark2-1)_d7(-)
speed(dark2-2)_d7(-)
speed(dark2-3)_d7(-)
speed(dark2-4)_d7(-)
speed(blue1)_d7(-)
speed(blue2)_d7(-)
speed(dark2-1)_d7(+)
speed(dark2-2)_d7(+)
speed(dark2-3)_d7(+)
speed(dark2-4)_d7(+)
speed(blue1)_d7(+)
speed(blue2)_d7(+)
speed(dark1-1)_d12(-)
speed(dark1-2)_d12(-)
speed(green1)_d12(-)
speed(green2)_d12(-)
speed(dark2-1)_d12(-)
speed(dark2-2)_d12(-)
speed(dark2-3)_d12(-)
speed(dark2-4)_d12(-)
speed(blue1)_d12(-)
speed(blue2)_d12(-)
speed(dark1-1)_d12(+)
speed(dark1-2)_d12(+)
speed(green1)_d12(+)
speed(green2)_d12(+)
speed(dark2-1)_d12(+)
speed(dark2-2)_d12(+)
speed(dark2-3)_d12(+)
speed(dark2-4)_d12(+)
speed(blue1)_d12(+)
speed(blue2)_d12(+)
resting_d7(+)
resting_d7(-)
resting_d12(+)
resting_d12(-)
anxiety_d7
anxiety_d12
LB(total)_d7(-)
LB(total)_d12(-)
LB(ratio)_d7(-)
LB(ratio)_d12(-)
LB(total)_d7(+)
LB(total)_d12(+)
phototaxis_d7(-)
phototaxis_d12(-)
phototaxis_d7(+)
thermotaxis
scrunching
NS(rate)(+)
NS(strength)(+)
NS(strength)(-)
Acephate
Chlorpyrifos
Diazinon
Dichlorvos
Malathion
Parathion
Profenofos
Aldicarb
Physostigmine
Anatoxin-A
Nicotine
Muscarine
Bethanechol
Buspirone
Mianserin
Fluoxetine
Sertraline
MDL-12,330A
LRE-1
Colchicine
Nocodazole
Cytochalasin D
Latrunculin A
Anandamide
WIN 55 212-2
L-buthionine sulfoxime
Rotenone
L-ascorbic acid
D-glucitol
(B)
Regenerating
lethality_d7
lethality_d12
shape_d7
shape_d12
stickiness_d7
stickiness_d12
speed(dark1-2)_d7(-)
speed(green1)_d7(-)
speed(green2)_d7(-)
speed(dark2-1)_d7(-)
speed(dark2-2)_d7(-)
speed(dark2-3)_d7(-)
speed(dark2-4)_d7(-)
speed(blue1)_d7(-)
speed(blue2)_d7(-)
speed(dark2-1)_d7(+)
speed(dark2-2)_d7(+)
speed(dark2-3)_d7(+)
speed(dark2-4)_d7(+)
speed(blue1)_d7(+)
speed(blue2)_d7(+)
speed(dark1-1)_d12(-)
speed(dark1-2)_d12(-)
speed(green1)_d12(-)
speed(green2)_d12(-)
speed(dark2-1)_d12(-)
speed(dark2-2)_d12(-)
speed(dark2-3)_d12(-)
speed(dark2-4)_d12(-)
speed(blue1)_d12(-)
speed(blue2)_d12(-)
speed(dark1-1)_d12(+)
speed(dark1-2)_d12(+)
speed(green1)_d12(+)
speed(green2)_d12(+)
speed(dark2-1)_d12(+)
speed(dark2-2)_d12(+)
speed(dark2-3)_d12(+)
speed(dark2-4)_d12(+)
speed(blue1)_d12(+)
speed(blue2)_d12(+)
resting_d7(+)
resting_d7(-)
resting_d12(+)
resting_d12(-)
anxiety_d7
anxiety_d12
LB(total)_d7(-)
LB(total)_d12(-)
LB(ratio)_d7(-)
LB(ratio)_d12(-)
LB(total)_d7(+)
LB(total)_d12(+)
phototaxis_d7(-)
phototaxis_d12(-)
phototaxis_d7(+)
thermotaxis
scrunching
NS(rate)(+)
NS(strength)(+)
NS(strength)(-)
eye

**Supplementary Figure 7. Heatmaps comparing the benchmark concentrations (BMCs) for tested chemicals in adult (A) and regenerating (B) planarians.** The first row shows the highest tested concentration. For endpoints that can have effects in both directions, the BMCs are separated by either the positive (+) or (-) direction.

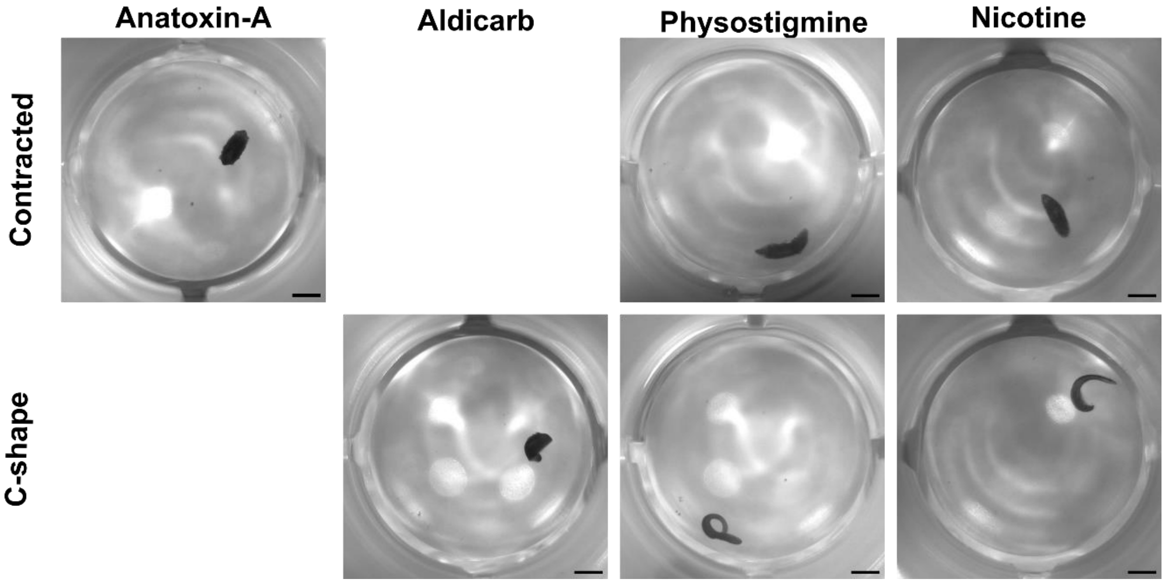


**Supplementary Figure 8. Examples of contracted and c-shape body shapes in cholinergic drugs.** Examples are of adult planarians exposed to the highest test concentration of the specified chemicals. Scale bar: 1 mm.
